# Genetic evidence for functional diversification of gram-negative intermembrane phospholipid transporters

**DOI:** 10.1101/2023.06.21.545913

**Authors:** Ashutosh K. Rai, Katsuhiro Sawasato, Haley C. Bennett, Anastasiia Kozlova, Genevieve C. Sparagna, Mikhail Bogdanov, Angela M. Mitchell

## Abstract

The outer membrane of Gram-negative bacteria is a barrier to chemical and physical stress. Phospholipid transport between the inner and outer membranes has been an area of intense investigation and, in *E. coli* K-12, it has recently been shown to be mediated by YhdP, TamB, and YdbH, which are suggested to provide hydrophobic channels for phospholipid diffusion, with YhdP and TamB playing the major roles. However, YhdP and TamB have different phenotypes suggesting distinct functions. We investigated these functions using synthetic cold sensitivity (at 30 °C) caused by deletion of *yhdP* and *fadR*, a transcriptional regulator controlling fatty acid degradation and unsaturated fatty acid production, but not by Δ*tamB* Δ*fadR* or Δ*ydbH* Δ*fadR*,. Deletion of *tamB* suppresses the Δ*yhdP* Δ*fadR* cold sensitivity suggesting this phenotype is related to phospholipid transport. The Δ*yhdP* Δ*fadR* strain shows a greater increase in cardiolipin upon transfer to the non-permissive temperature and genetically lowering cardiolipin levels can suppress cold sensitivity. These data also reveal a qualitative difference between cardiolipin synthases in *E. coli*, as deletion of *clsA and clsC* suppresses cold sensitivity but deletion of *clsB* does not despite lower cardiolipin levels. In addition to increased cardiolipin, increased fatty acid saturation is necessary for cold sensitivity and lowering this level genetically or through supplementation of oleic acid suppresses the cold sensitivity of the Δ*yhdP* Δ*fadR* strain. Although indirect effects are possible, we favor the parsimonious hypothesis that YhdP and TamB have differential substrate transport preferences, most likely with YhdP preferentially transporting more saturated phospholipids and TamB preferentially transporting more unsaturated phospholipids. We envision cardiolipin contributing to this transport preference by sterically clogging TamB-mediated transport of saturated phospholipids. Thus, our data provide a potential mechanism for independent control of the phospholipid composition of the inner and outer membranes in response to changing conditions.

**Author Summary:** Gram-negative bacteria possess a highly impermeable outer membrane, which protects against environmental stress and antibiotics. Outer membrane phospholipid transport remained mysterious until YhdP, TamB, and YdbH were recently implicated in phospholipid transport between the inner and outer membranes of *E. coli*. Similar roles for YhdP and/or TamB have been suggested in both closely and distantly related gram-negative bacteria. Here, given the transporters’ apparent partial redundancy, we investigated functional differentiation between YhdP and TamB. Our data suggest YhdP and TamB have differential involvement with fatty acid and phospholipid metabolism. In fact, transport of higher than normal levels of cardiolipin and saturated phospholipids in the absence of YhdP and presence of TamB at a non-permissive temperature is lethal. We suggest a model where the functions of YhdP and TamB are distinguished by phospholipid transport preference with YhdP preferentially transporting more saturated phospholipids and TamB more unsaturated phospholipids. Cardiolipin headgroup specificity may contribute transport inhibition due to its bulky nature inhibiting the passage of other phospholipids. Diversification of function between YhdP and TamB provides a mechanism for regulation of phospholipid composition, and possibly the mechanical strength and permeability of the outer membrane, and so the cell’s intrinsic antibiotic resistance, in changing environmental conditions.

## Introduction

The gram-negative bacterial cell envelope has an outer membrane (OM) that sits outside the aqueous periplasm and peptidoglycan cell wall. The OM provides a barrier against various environmental stresses including toxic molecules such as antibiotics and osmotic pressure (1-6). Unlike the inner membrane (IM), a phospholipid bilayer, the OM is largely composed of phospholipids (mainly phosphatidylethanolamine (PE) (7)) in its inner leaflet and LPS (lipopolysaccharide) in its outer leaflet (2, 8). However, both membranes are asymmetric as the IM has different phospholipid compositions in its inner versus outer leaflets (9). Outer membrane proteins (OMPs) form a network across the cell surface interspersed with phase separated LPS patches (10). Lipoproteins are generally anchored in the inner (i.e., periplasmic) leaflet of the OM (11).

OM components are synthesized in the cytoplasm or IM and transported to the OM. OMP (12), lipoprotein (11), and LPS (13) transport pathways are well defined. However, until recently, intermembrane phospholipid transport (between the IM and OM), especially anterograde transport from the IM to the OM, has remained very poorly understood. Phospholipids are synthesized at the IM’s inner leaflet (14) and rapid, bidirectional intermembrane phospholipid transport occurs (15), even without ATP (16). Phospholipid transport has a relaxed specificity, allowing transport of non-native lipids (17, 18); however, intermembrane phospholipid composition differences are maintained (7, 19-22). For instance, the OM is enriched for PE and its saturated species compared to the IM outer leaflet (7, 9, 20-22); however, phosphatidylglycerol (PG) and cardiolipin (CL) can be evenly distributed between the IM and OM when the cellular level of PE is greatly reduced (17), suggesting a maintenance of membrane charge balance. This headgroup redistribution is accompanied by a concomitant fatty acid redistribution evidenced by an accumulation of palmitic acid (C16:0) and cyclopropane derivatives of palmitoleic acid (C16:1) and *cis*-vaccenic acid (C18:1). In contrast, the 1,2-diglyceride that accumulates in mutants lacking DgkA (diglyceride kinase) predominantly associates with the IM (23), suggesting existence of a mechanism of discrimination of lipid transfer to the OM that does not recognize diglyceride molecules and contributes to the diversification of distribution of polar and acyl groups between the IM and OM. This establishment and maintenance of defined lipid topography between the IM and the OM must involve the coordinated biosynthesis and balanced bidirectional trafficking of specific phospholipids across the IM and from the IM to the OM (24).

It has been demonstrated that anterograde phospholipid transport is mediated by a high-flux, diffusive (i.e., concentration gradient, not energy dependent) system in *Escherichia coli* (25). A retrograde trafficking pathway, the Mla system, removes mislocalized phospholipids from the OM outer leaflet and returns them to the IM (26). A gain-of-function allele, *mlaA\** (27, 28), opens a channel allowing phospholipids to mislocalize to the cell surface by flowing from the OM’s inner to outer leaflet (27, 29), resulting in a PldA (OM phospholipase A)-mediated attempt to reestablish OM asymmetry by increasing LPS production, OM vesiculation, and increased flow of phospholipids to the OM (27, 28). When nutrients are depleted, inhibiting phospholipid production, cells lyse due to loss of IM integrity (16, 18). Single cell imaging of *mlaA\** cells confirmed an aberrant lipid flow from the IM to the OM at a fast rate. Loss of YhdP, an inner membrane protein, could significantly slow phospholipid flow to the OM resulting in loss of OM integrity before IM rupture (25), suggesting YhdP might play a role in intermembrane phospholipid transport.

Recently, YhdP and two homologs, TamB and YdbH, were demonstrated to be intermembrane phospholipid transporters (30, 31). Ruiz, *et al*. investigated AsmA family proteins and found Δ*yhdP* Δ*tamB* mutants showed synthetic OM permeability and stationary phase lysis (30). Moreover, when combined, Δ*yhdP*, Δ*tamB*, and Δ*ydbH* are synthetically lethal. Genetic interactions of these mutants with *mlaA* and *pldA* show their involvement in OM phospholipid homeostasis. Predicted structures of YhdP, TamB, and YdbH (32, 33), as well as a partial crystal structure of TamB (34), have a β-taco fold domain with a hydrophobic pocket, resembling eukaryotic lipid transporters (35-38). These structural findings are another clue that suggests YhdP, TamB, and YdbH are involved in phospholipid transport. Douglass, *et al.* confirmed the synthetic lethality of Δ*yhdP*, Δ*tamB*, and Δ*ydbH* and demonstrated a Δ*tamB* Δ*ydbH* mutant with reduced *yhdP* expression had decreased amounts of OM phospholipids, directly demonstrating their involvement in intermembrane phospholipid transport or its regulation (31). Structural studies have shown that YhdP is long enough to span the periplasmic space and molecular dynamics indicate that the C-terminus of YhdP can insert into the OM to allow phospholipid transfer between the membranes (39). This study also directly demonstrates that a phosphate containing substrate, putatively assigned to be phospholipids, can be crosslinked to the hydrophobic groove of YhdP (39). The role of YhdP, TamB, and YdbH in phospholipid transport may be widely conserved as recent evidence implicates TamB in anterograde phospholipid transport in *Veillonella parvula*, a diderm Fermicute (40) and YhdP, TamB, YdbH, and PA4735 in intermembrane phospholipid transport in *Pseudomonas aeruginosa* (41).

Nevertheless, why there are three separate intermembrane phospholipid transport proteins and the functional interactions between these proteins remains unanswered, although it is clear the proteins are not fully redundant or functionally equivalent. The conditional expression of *yhdP* alone, without the presence of *ydbH* and *tamB*, fully complements neither growth phenotypes nor a normal level of OM phospholipids, suggesting transport function is still impaired in this strain (31). In addition, YdbH seems to play a more minor role than YhdP and TamB (30, 31) and the screen identifying Δ*yhdP* as slowing *mlaA\**-dependent lysis did not identify Δ*tamB* or Δ*ydbH* (25). Moreover, we previously identified a role for YhdP in stationary phase SDS (sodium dodecyl sulfate) resistance and modulating cyclic enterobacterial common antigen activity (42, 43) not shared by TamB and YdbH (**Fig. S1AB**) (30, 31, 42, 43). Similarly, TamB, in conjunction with TamA, has been suggested to play a role in OM insertion of some “complicated” β-barrel OMPs such as autotransporters and usher proteins (34, 44-46).

Here, we investigated the differentiation of YhdP and TamB function and identified synthetic cold sensitivity in a strain with Δ*yhdP* and Δ*fadR*, a transcriptional regulator of fatty acid biosynthesis and degradation, acting as a transcriptional switch between these pathways in response to acyl-CoA (47). FadR is necessary for normal levels of unsaturated fatty acids to be synthesized (48). The cold sensitivity was unique to the Δ*yhdP* Δ*fadR* strain and was suppressed by loss of TamB, demonstrating that the sensitivity is related to phospholipid transport. We found that the Δ*yhdP* Δ*fadR* mutant had a greater increase in levels of a specific phospholipid, cardiolipin (CL) during growth at the non-permissive temperature, and decreasing CL suppressed cold sensitivity. Furthermore, increasing unsaturated fatty acid levels suppressed cold sensitivity. Overall, our data are consistent with a model where TamB and YhdP transport functions are differentiated based on phospholipid fatty acid saturation state, with TamB transporting more unsaturated phospholipids and YhdP transporting more saturated phospholipids, although indirect effects may contribute to the phenotype. Phospholipid species or head group can also contribute to transport specificity, since higher levels of dianionic CL are necessary for the cold sensitivity phenotype, potentially due to functional inhibition of TamB by clogging of its hydrophobic groove.

## Results

### Loss of FadR and YhdP results in cold sensitivity suppressed by loss of TamB

To investigate YhdP-TamB function differentiation, we created a Δ*yhdP* Δ*fadR* strain, as FadR is a major regulator of lipid homeostasis (47). A deletion strain of *fadR* alone is expected to have decreased fatty acid synthesis, increased fatty acid degradation, and increased fatty acid saturation (47, 48). Surprisingly, Δ*yhdP* Δ*fadR* cultures grown at 30 °C lagged for more than 8 hours, followed by highly variable growth, suggesting suppressor outgrowth (**Fig.1A**). The growth defect was decreased at 37 °C and completely absent at 42 °C. To confirm this cold sensitivity, we estimated efficiency of plating (EOP) and confirmed the severe cold intolerance of the Δ*yhdP* Δ*fadR* strain (5 logs at 30 °C and 3 logs at 37 °C) (**Fig. 1B**). Although, enterobacterial common antigen is necessary for Δ*yhdP*’s OM permeability phenotypes (43), the Δ*yhdP* Δ*fadR* strain’s cold sensitivity was unchanged when enterobacterial common antigen was not present (Δ*wecA*) (49) (**Fig. 1C**).

**Figure 1.**
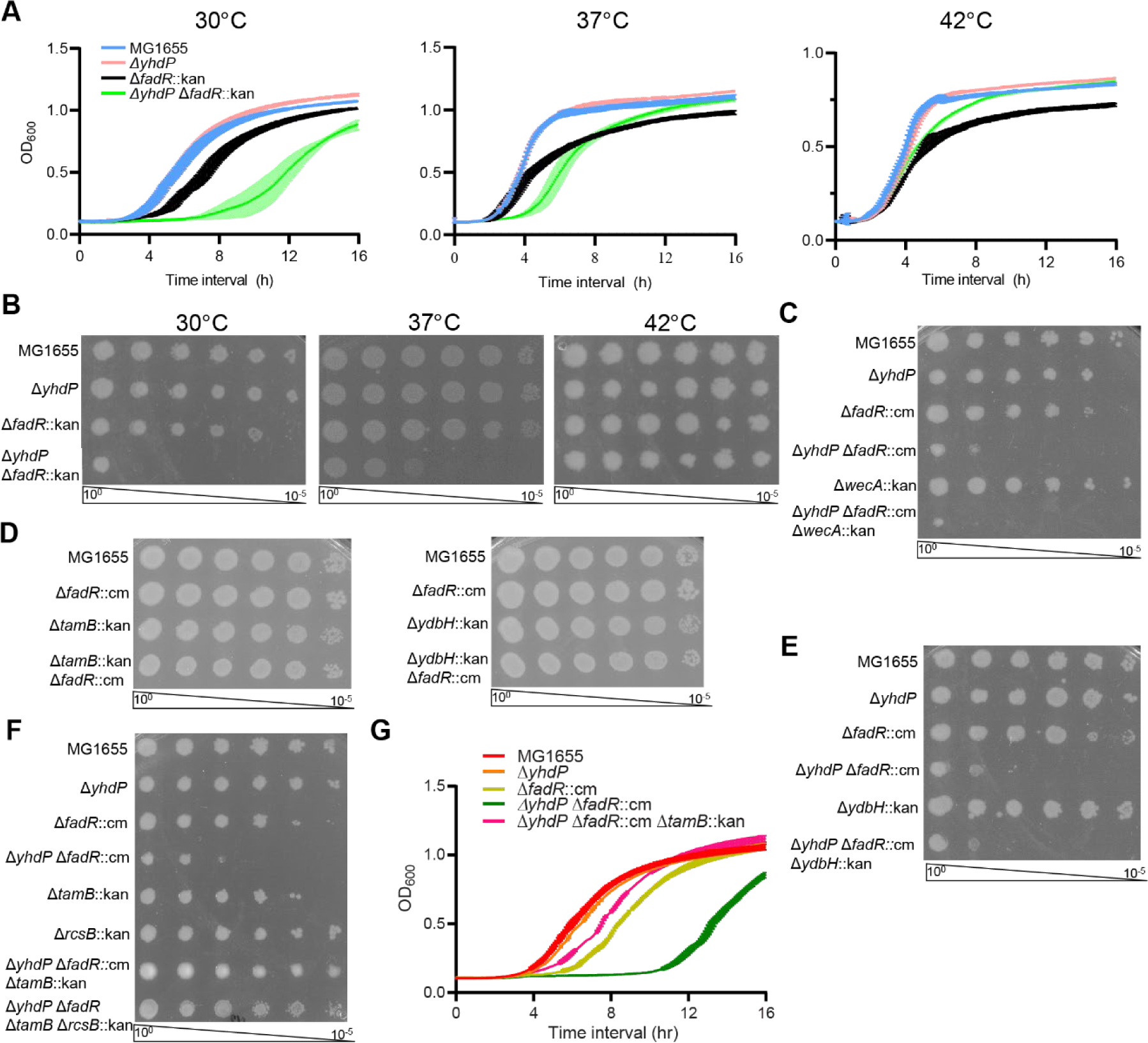
Deletion of *yhdP* and *fadR* causes synthetic cold sensitivity for which TamB is necessary. **(A)** The Δ*yhdP* Δ*fadR* strain had impaired growth at lower temperatures. **(B)** EOPs were performed to confirm the growth defect of the Δ*yhdP* Δ*fadR* strain. **(C-F)** EOPs were performed at 30 °C to assess cold sensitivity. **(C)** The growth of the Δ*yhdP* Δ*fadR* strain was not affected by the loss of ECA (Δ*wecA*). **(D)** Neither a Δ*tamB* Δ*fadR* strain nor a Δ*ydbH* Δ*fadR* strain exhibited cold sensitivity. **(E)** Deletion of *ydbH* did not suppress cold sensitivity in the Δ*yhdP* Δ*fadR* strain. **(F)** Deletion of *tamB* completely suppressed cold sensitivity in the Δ*yhdP* Δ*fadR* strain and the Rcs stress response was not necessary for this suppression. **(G)** Growth curves were performed at 30 °C. The Δ*yhdP* Δ*fadR* Δ*tamB* strain grew well, indicating suppression was unlikely to result from slow growth. Quantitative data are shown as the mean of three biological replicates ± the SEM. Images are representative of three independent experiments.

We hypothesized the Δ*yhdP* Δ*fadR* strain’s cold sensitivity was due to impairment of phospholipid transport and expected Δ*tamB* and/or Δ*ydbH* to show similar synthetic phenotypes with Δ*fadR*. However, Δ*fadR* Δ*tamB* and Δ*fadR* Δ*ydbH* strains had no growth defects (**Fig 1D**). Although Δ*ydbH* in the Δ*yhdP* Δ*fadR* strain did not alter cold sensitivity (**Fig. 1E**), Δ*tamB* in the Δ*yhdP* Δ*fadR* background completely suppressed the cold sensitivity (**Fig. 1F**). Consistent with previous observations for Δ*yhdP* Δ*tamB* (30), the Δ*yhdP* Δ*fadR* Δ*tamB* strain was mucoid due to Rcs (regulator of capsule synthesis) stress response activation and colanic acid capsule production. Deleting *rcsB*, an Rcs response regulator, prevented mucoidy but did not affect suppression (**Fig. 1F**). We observed only minimal growth phenotypes at 30 °C in the Δ*yhdP* Δ*fadR* Δ*tamB* strain (**Fig. 1G**), similar to those of Δ*yhdP* Δ*tamB* (**Fig. S1C**). Thus, although Δ*fadR* does not suppress Δ*yhdP* Δ*tamB* phenotypes, these data make slow growth an unlikely suppression mechanism. Together, these data suggest the Δ*yhdP* Δ*fadR* strain has impaired phospholipid transport leading synthetic cold sensitivity, which can be relieved by deletion of Δ*tamB*, shifting phospholipid transport to YdbH or causing some regulatory change.

We next looked for envelope homeostasis alterations in the Δ*yhdP* Δ*fadR* strain. To identify cell lysis and/or severe OM permeability defects at 30 °C, we assayed CPRG (chlorophenol red-β-D-galactopyranoside) processing. CPRG must contact LacZ in the cytoplasm or supernatant to release chlorophenol red (50). Compared to wild type or single mutants, the Δ*yhdP* Δ*fadR* strain had increased CPRG processing (**Fig. S2A**), demonstrating lysis or increased envelope permeability. When we assayed resistance to molecules excluded by the OM (1, 42, 43, 51, 52) at 37 °C, a semi-permissive temperature, the Δ*yhdP* Δ*fadR* strain showed sensitivity to vancomycin, a large scaffold antibiotic, and SDS EDTA compared to single mutants (**Fig. S2B**). However, with bacitracin treatment or EDTA treatment alone, the Δ*yhdP* Δ*fadR* strain phenocopied the Δ*fadR* single deletion (**Fig. S2CD**). LPS levels were not altered in the Δ*yhdP*, Δ*fadR*, Δ*yhdP* Δ*fadR* or Δ*yhdP* Δ*fadR* Δ*tamB* strains (**Fig. S2E**). OM asymmetry mutations (i.e., Δ*pldA*, Δ*mlaA*) (26-28, 53) did not have suppressive or synthetic effects on cold sensitivity in the Δ*yhdP* Δ*fadR* strain (**Fig. S3AB**), cold sensitivity is not due to OM asymmetry or the loss thereof. Overall, the changes in OM permeability did not seem sufficient to cause lethality at lower temperature, suggesting the phospholipid transport disruption affects both IM and OM integrity.

### Lowering cardiolipin suppresses cold sensitivity

We investigated phospholipid composition of the wild-type, Δ*yhdP*, Δ*fadR*, and Δ*yhdP* Δ*fadR* strains grown at the permissive temperature (42 °C) to log phase then downshifted to 30 °C. *E. coli* phospholipid composition is generally 75% phosphatidylethanolamine (PE), 20% phosphatidylglycerol (PG), and 5% cardiolipin (CL) with CL increasing in stationary phase (14, 54). Of the three CL synthases, ClsA and ClsB synthesize CL from two PG molecules, while ClsC synthesizes CL from one PG and one PE molecule, so CL and PG levels are generally reciprocally regulated (55-58). For all strains, the levels of PE were similar (**Fig. 2A**, **Fig. S4A**). However, the Δ*yhdP* Δ*fadR* strain had increased CL and concomitant decreased PG compared to the wild-type strain and compared to individual Δ*yhdP* and Δ*fadR* mutants (**Fig. 2B**). PgsA synthesizes phosphatidylglycerol-phosphate and is the first enzyme differentiating PG and PE biosynthesis. PgsA depletion would decrease both PG and CL levels, reversing increased CL levels and exacerbating decreased PG levels in the Δ*yhdP* Δ*fadR* strain. We tested the effect of PgsA depletion in a strain background (Δ*rcsF* Δ*lpp*) where *pgsA* is non-essential (59-62). When *pgsA* expression was repressed (glucose panel), the Δ*yhdP* Δ*fadR* cold sensitivity was partially suppressed (**Fig. 2C**), demonstrating that decreased PG does not cause cold sensitivity and suggesting increased CL may be necessary for cold sensitivity. The partial suppression may be due to incomplete depletion of PG and CL or the adaptive response of a cell containing only PE in its membrane. Induction of *pgsA* (arabinose panel) reversed the suppression. The cold sensitivity of the Δ*yhdP* Δ*fadR* strain was lost when CL was fully depleted due to deletion of the genes for all cardiolipin syntheses (Δ*clsABC*) or of the gene for the primary cardiolipin synthase (Δ*clsA*) responsible for CL synthesis in exponential phase (**Fig. 2ADE, Fig. S4B**). A decrease in LPS levels is not responsible for the suppression as LPS levels did not change in the Δ*yhdP* Δ*fadR* Δ*clsA* strain (Δ*yhdP* Δ*fadR* Δ*tamB*). Δ*clsB* caused a smaller decrease in CL levels, while Δ*clsC* did not cause a decrease (**Fig. 2AB**, **Fig. 4B**). Surprisingly, however, Δ*clsC* completely suppressed the cold sensitivity, while Δ*clsB* did not (**Fig. 2FG**). Thus, increased CL is necessary for cold sensitivity but CL levels alone cannot explain the phenotype: another factor, such as a specific molecular form (e.g., acyl chain length, saturation state, symmetry of acyl chain arrangement) or localization (e.g., poles vs. midcell or inner vs. outer leaflet) of CL, is required.

**Figure 2.**
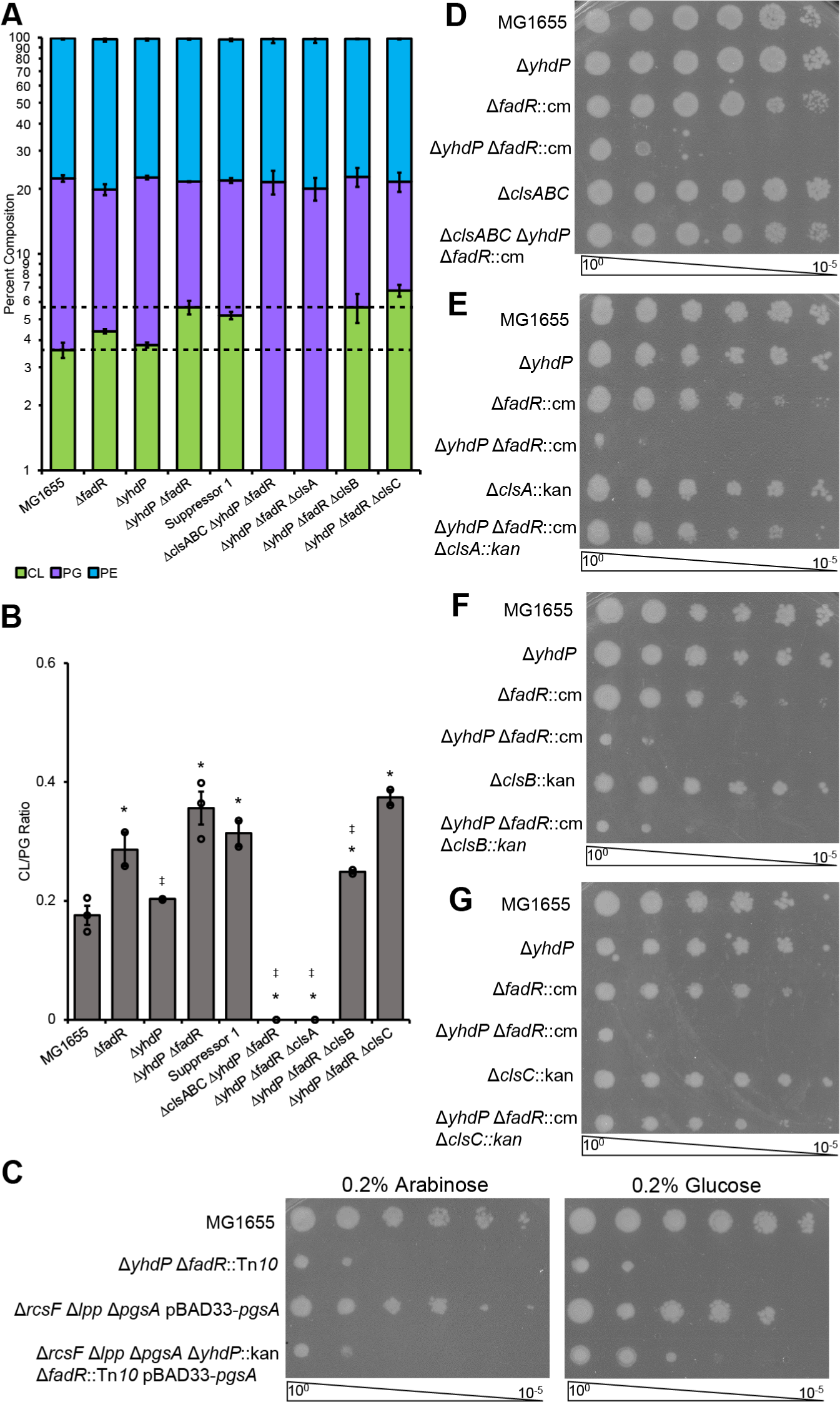
Lowering cardiolipin levels can suppress cold sensitivity of the Δ*yhdP* Δ*fadR* strain. **(A-B)** Thin layer chromatography (TLC) of lipid extracts was performed for the indicated cultures grown to log phase at 42 °C then transferred to 30 °C for 2 hours before analysis. **(A)** Relative phospholipid levels were calculated. The Δ*yhdP* Δ*fadR* strain shows increased levels of CL and a concomitant decrease in PG levels, increasing the ratio of CL to PG (B). Data are averages of two to three biological replicates ± the SEM. For ratios individual data points are indicated with open circles. * p<0.05 vs. MG1655 by Student’s T-test; ‡ p<0.05 vs. the Δ*yhdP* Δ*fadR* strain by Student’s T-test. **(C-G)** Growth effects of altered phospholipid levels were investigated by EOP. Images are representative of three independent experiments. **(C)** PgsA depletion strains in a background where *pgsA* is non-essential were used to assay the effect of lowering PG and CL levels on cold sensitivity. Arabinose induces expression of *pgsA* while glucose represses. Cultures were induced or repressed for 4-5 generations before as well as during the EOP. Lowering PG and CL levels partially suppressed cold sensitivity. **(D)** Deletion of the three CL synthases, *clsA*, *clsB*, and *clsC*, suppressed cold sensitivity of the Δ*yhdP* Δ*fadR* strain. (E) Deletion of *clsA* also suppressed cold sensitivity. **(F)** Deletion of *clsB* does not suppress cold sensitivity, while deletion of *clsC* (F) does.

### Increased *fabA* expression suppresses cold sensitivity

To identify other factors involved in the cold sensitivity, we applied a forward genetic approach by isolating spontaneous suppressor mutants capable of growing at 30 °C and identified a suppressor mutant (Suppressor 1) that restored growth at 30 °C (**Fig. 3A**), decreased CPRG processing (**Fig. S2A**), and had very similar CL levels (**Fig. 2AB**, **Fig. S4A**) and LPS levels (**Fig. S2**) to the Δ*yhdP* Δ*fadR* strain. We identified a point mutation in the *fabA* promoter region in Suppressor 1 (**Fig. 3B**) and confirmed this mutation was sufficient for suppression (**Fig. 3C**). FabA is a dehydratase/isomerase that introduces *cis* unsaturation into fatty acids (14, 63) and *fabA* is expressed from two promoters controlled by FadR and FabR (64, 65). FadR activates *fabA* expression, while FabR represses. Thus, *fadR* deletion would be expected to lower *fabA* expression. The suppressor mutation was located in the *fabA* promoter FabR binding site (**Fig. 3B**) and we hypothesized this mutation would increase *fabA* expression. *fabA* mRNA levels were more than 4-fold lower in the Δ*yhdP* Δ*fadR* strain than the wild-type strain (**Fig. 3D**). While lower than wild type, the suppressor mutant increased *fabA* mRNA 1.6-fold over the Δ*yhdP* Δ*fadR* strain.

**Figure 3.**
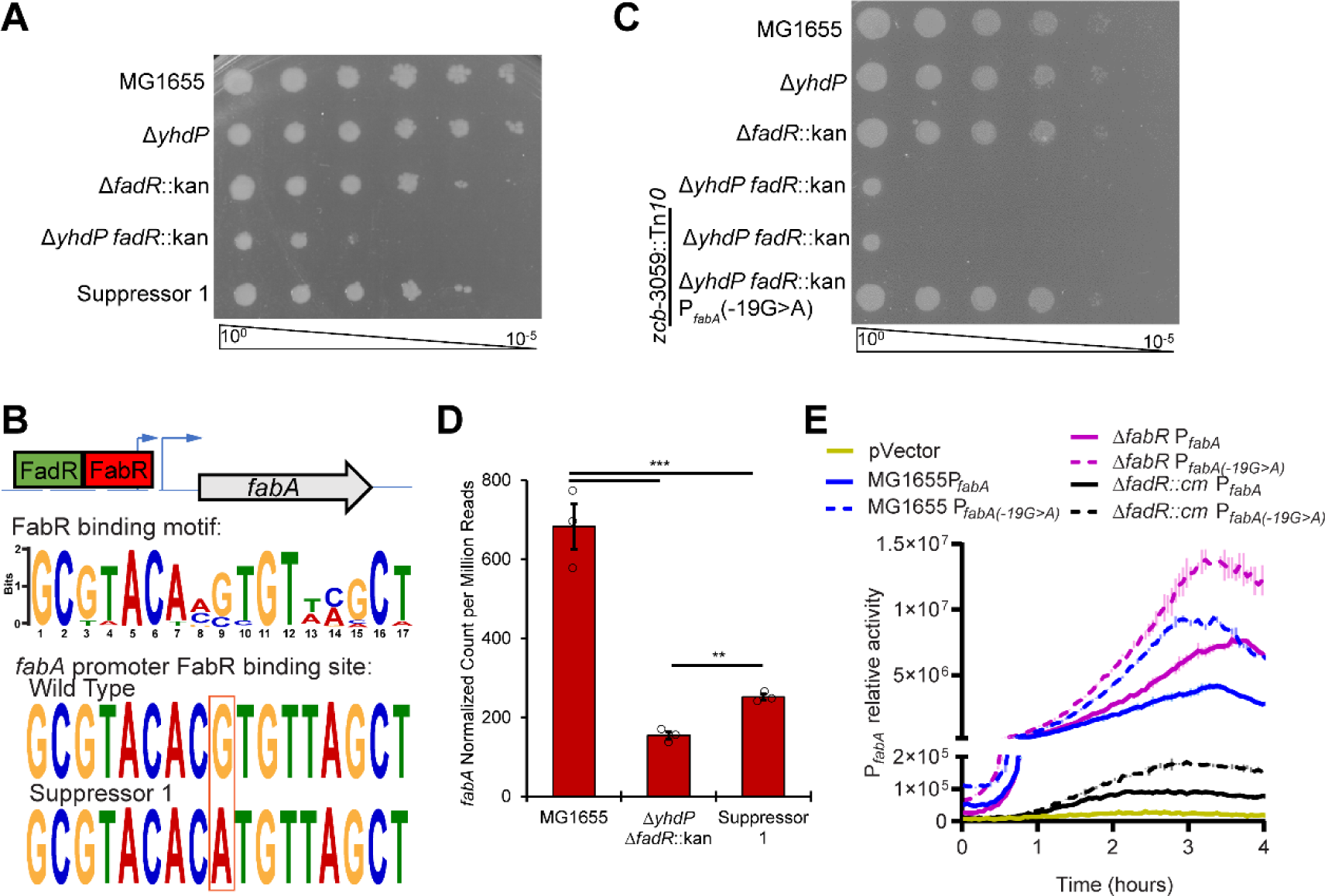
Cold sensitivity can be suppressed by increasing *fabA* expression. **(A)** The cold sensitivity of a spontaneous suppressor of Δ*yhdP* Δ*fadR* cold sensitivity was tested by EOP. The suppressor returned growth to the level of the Δ*fadR* mutant. **(B)** A diagram of the promoter region of *fabA* is shown with its transcriptional start sites (blue arrows) and regulator binding sites (boxes). FadR acts as an activator while FabR acts as a repressor. The motif of the FabR binding site, generated using MEME, is shown with the sequences of the parent strain and of Suppressor 1. **(C)** The P*_fabA_*(-19G>A) mutation was linked to a Tn*10* and transferred to the Δ*yhdP* Δ*fadR* strain by transduction. The Δ*yhdP* Δ*fadR* strain with P*_fabA_*(-19G>A) was not cold sensitivity, confirming suppression. **(D)** *fabA* RNA levels are shown. Bars are the mean of three biological replicates ± the SEM. Individual data points are shown as open circles. ** p<0.005, *** p<0.0005 by quasi-linear F-test. Data are from the RNA-seq experiment in Figure 5 and *fabA* data are also shown in Figure 5F. **(E)** Luciferase reporters of wild-type and mutant *fabA* promoter activity were constructed. The mutation caused similar fold increases in P*_fabA_* activity in all strain backgrounds, indicating the activity of the second *fabA* promoter and not FabR binding were likely affected by the suppressor mutation. Data are luminescence relative to OD_600_ and are shown as the average of two biological replicates ± the SEM.

We constructed luciferase reporter plasmids with the wild-type *fabA* promoter region or the *fabA* promoter region containing the suppressor mutation. The reporter with the mutated promoter had higher activity in a wild-type, Δ*fadR,* and Δ*fabR* background (**Fig. 3E**). *fabA* transcription occurs from a promoter downstream of the FabR binding site (65-67). However, when *fadR* is deleted, transcription shifts to the FabR-regulated promoter within the FabR binding region (65-67). Thus, our data indicate the Suppressor 1 mutation regulates the constitutive activity of the promoter located in the FabR binding site rather than the affinity of FabR binding. To confirm the effect of changing the second promoter’s activity, we tested the effect of Δ*fabR* on cold sensitivity in the Δ*yhdP* Δ*fadR* strain and observed partial suppression (**Fig. S5**). The smaller effect of Δ*fabR* compared to Suppressor 1 may result from other gene expression changes or to the smaller relative effect of Δ*fabR* on *fabA* expression (**Fig. 3E**). Overall, these data demonstrate increasing *fabA* expression, which is necessary for unsaturated fatty acid biosynthesis, rescues cold sensitivity in the Δ*yhdP* Δ*fadR* strain without changing CL levels.

### Increasing unsaturated fatty acids relieves cold sensitivity

Given the effect of *fabA* expression on the Δ*yhdP* Δ*fadR* cold sensitivity, we used liquid chromatography-electrospray ionization mass spectrometry (LC/MS) to characterize the strains’ phospholipid saturation state (**Dataset S1**, **Fig. 4A-C, Fig. S6**). Representative spectra demonstrating relative phospholipid composition are shown in **Fig. S7**. As expected, Δ*fadR* caused increased fully saturated and monounsaturated PE and PG with a concomitant decrease of diunsaturated PE (**Fig. 4AB, S6AB**). Similarly, the Δ*fadR* strain demonstrated increased monounsaturated CL with a concomitant decrease in triunsaturated CL (**Fig. 4C, S6C**). Compared to the Δ*fadR* strain, the Δ*yhdP* Δ*fadR* strain had slightly decreased saturation for all three phospholipids, while still displaying more phospholipid saturation than wild type or Δ*yhdP* strains (**Fig. 4A-C, S6C**). Suppressor 1 trended towards decreased saturation of all three phospholipids compared to the Δ*yhdP* Δ*fadR* strain (**Fig. 4A-C, S6C**). We wondered whether the suppression difference between Δ*clsC* and Δ*clsB* resulted from the CL saturation state. However, the Δ*yhdP* Δ*fadR* Δ*clsB* and Δ*yhdP* Δ*fadR* Δ*clsC* strains had very similar CL profiles (**Fig. 4C, S6C**), suggesting another qualitative difference affecting suppression, perhaps CL lateral or inter-leaflet distribution in the cell. These strains also showed similar PE and PG saturation profiles to their Δ*yhdP* Δ*fadR* parent (**Fig. 4AB, S6AB**).

**Figure 4.**
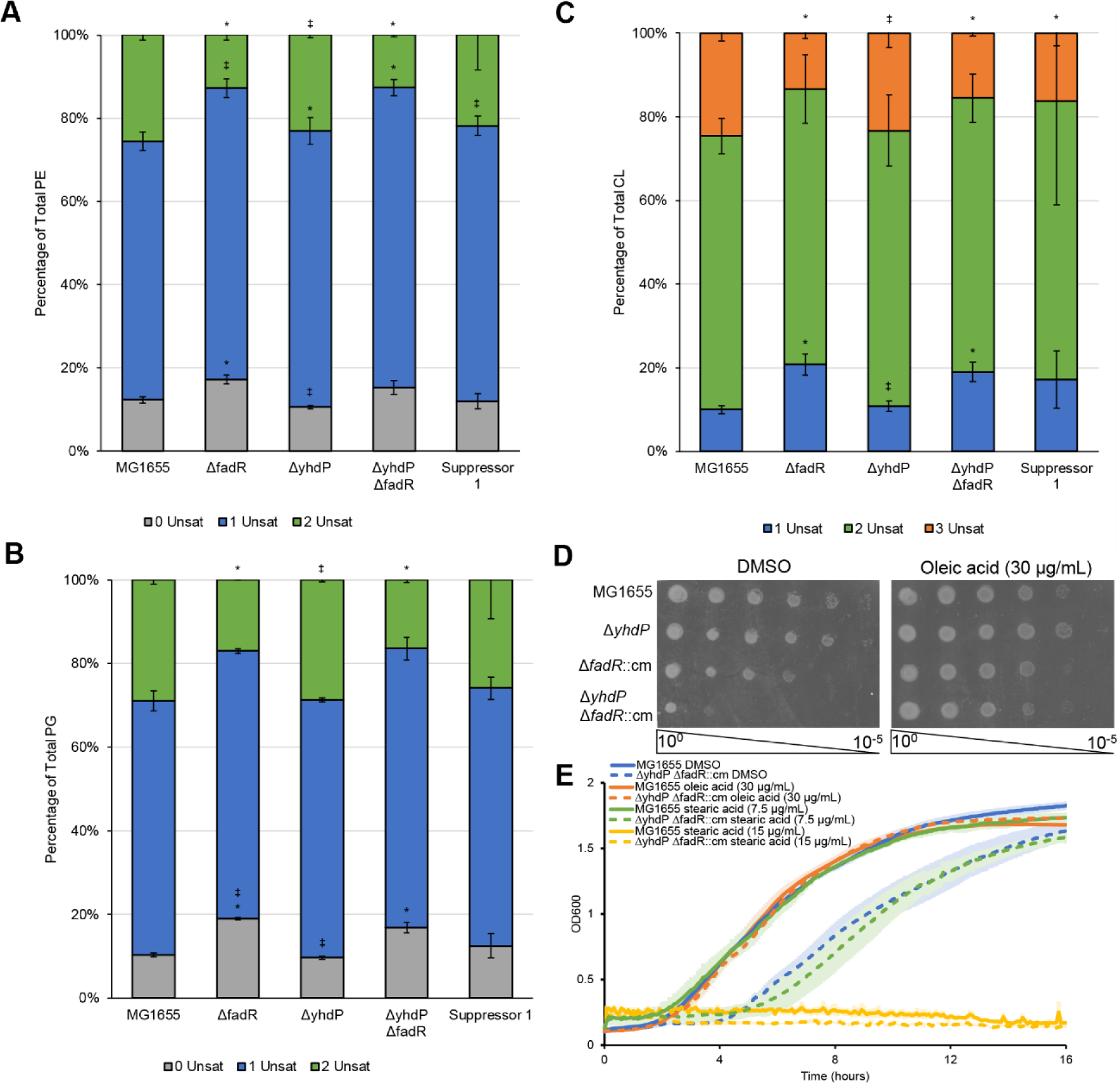
Decreasing fatty acid saturation suppresses Δ*yhdP* Δ*fadR* cold sensitivity. **(A-C)** Lipid were extracted from cells grown to OD_600_ 0.2 at 42 °C then shifted to 30 °C for 2 hours before harvest. LC/MS was performed on lipid extracts using absolute quantification and the percentage of each specific species of PE (A), PG (B), and CL (C) was calculated using the sum of all molecular species detected. Suppressor 1 trends towards decreased saturation compared to its parent strain (Δ*yhdP* Δ*fadR*). Data for phospholipids without unsaturations are shown in grey, with one unsaturation in blue, with two unsaturations in green, with three unsaturations in orange. Data are the average of three biological replicates ± the SEM. * p<0.05 vs. MG1655 by the Mann-Whitney test; ‡ p<0.05 vs. the Δ*yhdP* Δ*fadR* strain by the Mann-Whitney test. All molecular species are shown in Fig. S6. **(D)** EOPS were performed on media supplemented with oleic acid or a vehicle control. Oleic acid suppresses the cold sensitivity of the Δ*yhdP* Δ*fadR* strain. **(E)** Growth curves of the indicated strains were performed in the presence of oleic acid, stearic acid, or a vehicle control. Oleic acid suppressed the cold sensitivity of the Δ*yhdP* Δ*fadR* strain while stearic acid did not. Data are the average of three biological replicates ± the SEM.

We hypothesized increasing unsaturated fatty acids suppresses Δ*yhdP* Δ*fadR* cold sensitivity and performed EOPs for cold sensitivity on media containing an unsaturated fatty acid (oleic acid, C18:1 *cis*-9). Exogenous phospholipids can be taken up by *E. coli*, attached to acyl-CoA, and incorporated into phospholipids (68). Oleic acid addition suppressed the Δ*yhdP* Δ*fadR* strain’s cold sensitivity (**Fig. 4D**). Supplementing oleic acid increases unsaturated phospholipids and provides an exogenous fatty acid source. To differentiate between saturation state and fatty acid availability, we compared treatment with oleic acid and saturated stearic acid (C18:0). Only oleic acid, and not stearic acid, suppressed the Δ*yhdP* Δ*fadR* strain’s cold sensitivity (**Fig. 4E**), confirming increasing unsaturated fatty acids, not fatty acid availability suppresses.

### Suppression causes fatty acid metabolism and stress response alterations

In wild-type cells, *fabA* overexpression does not change phospholipid saturation levels, demonstrating the FabB enzyme is rate limiting for unsaturated fatty acid biosynthesis (69). We were intrigued that Suppressor 1 altered saturation without directly effecting *fabB* expression. We also wondered whether CL levels in the Δ*yhdP* Δ*fadR* strain are transcriptionally regulated. Thus, we investigated the transcriptional landscape of the wild type, the Δ*yhdP* Δ*fadR*, and Suppressor 1 strains after a 30-minute downshift of growth to 30 °C to determine: (i) whether the Δ*yhdP* Δ*fadR* strain had altered *cls* gene expression; (ii) whether Suppressor 1 had other transcriptional changes; and (iii) why increased *fabA* expression decreased saturation. All differentially expressed genes (>2-fold change, p<0.05) are listed in **Dataset S2**. Principle component analysis of these genes showed the closest relation between Δ*yhdP* Δ*fadR* strain and Suppressor 1, while the wild-type strain was more distantly related (**Fig. 5A**, **S8A**). Nevertheless, Suppressor 1 was more closely related to wild type than was the Δ*yhdP* Δ*fadR* strain. Most differentially regulated genes between Δ*yhdP* Δ*fadR* and wild type and Suppressor 1 and wild type were upregulated (**Fig. 5BC**, **S8BC**). Many of the most highly regulated genes are FadR-regulon members. The majority of differentially regulated genes in Suppressor 1 compared to Δ*yhdP* Δ*fadR* are downregulated (**Fig. 5D, S8D**). These genes are involved in many cellular pathways and likely reflect the decreased cellular stress (**Table S2**). Indeed, many external stress response genes are more enriched in Δ*yhdP* Δ*fadR* than in Suppressor 1 (**Fig. S9A**).

**Figure 5.**
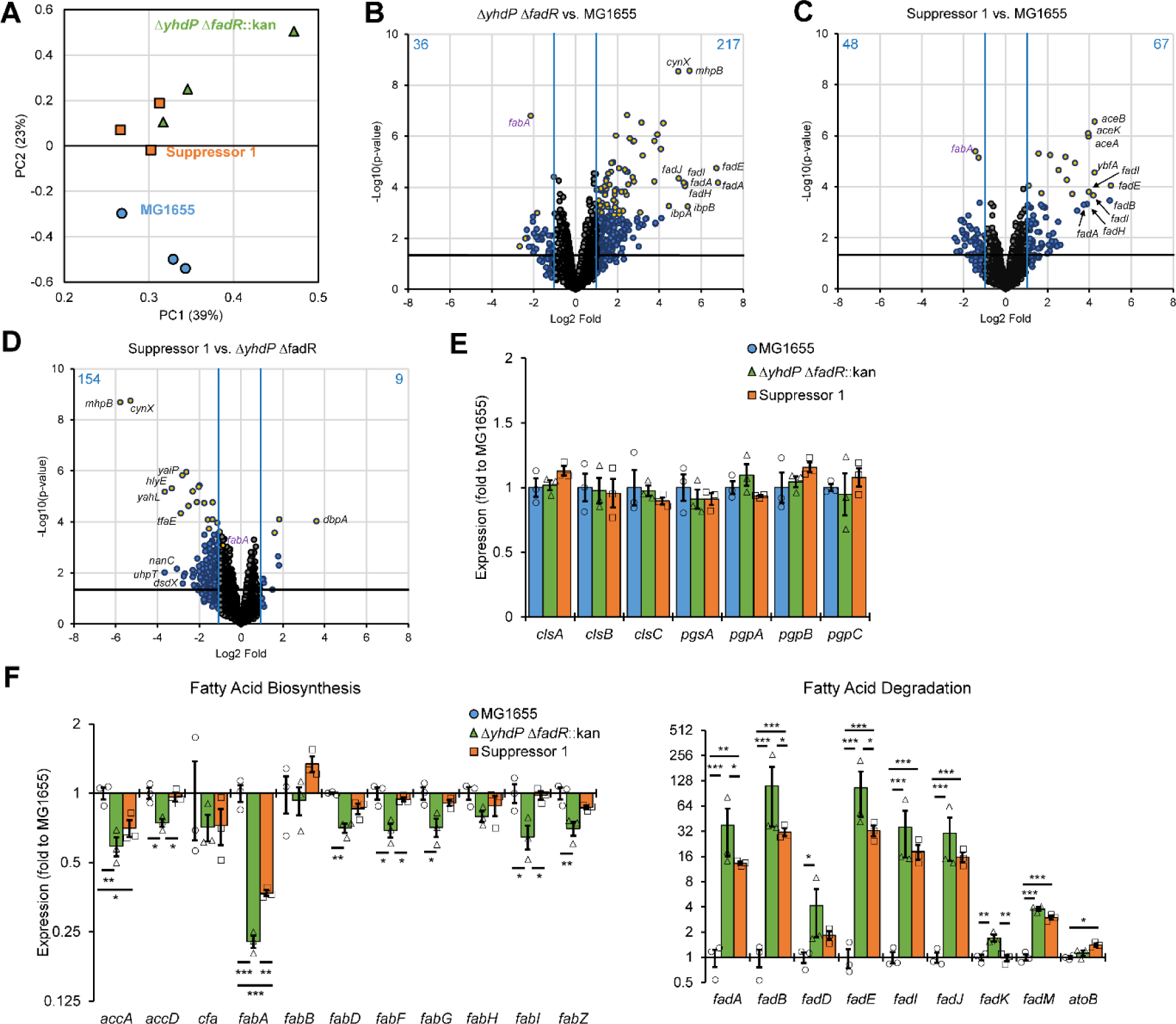
Transcriptional landscape of the Δ*yhdP* Δ*fadR* and suppressed strain. Three biological replicates of the indicated strains were grown at 42 °C to mid-log then transferred to 30 °C for 30 minutes before harvesting RNA and performing RNA-seq. Differential expression was calculated based on a greater than 2-fold change in expression and a p value of less than 0.05 by quasi-linear F-test. **(A)** PCA was performed on the expression of all genes differentially expressed between any groups of samples. Suppressor 1 and the Δ*yhdP* Δ*fadR* strain grouped closer together than to the wild-type strain, although Suppressor 1 was closer to wild type than the Δ*yhdP* Δ*fadR* strain. **(B-D)** Volcano plots show the average fold change and significance of expression changes between the Δ*yhdP* Δ*fadR* strain and wild-type MG1655 (B), Suppressor 1 and wild-type MG1655 (C), and Suppressor 1 and the Δ*yhdP* Δ*fadR* strain (D). Genes with changes greater than 2-fold are shown in blue, while genes with changes less than 2-fold are shown in grey. Genes with a q-value (false discovery rate) of less than 0.05 are shown in yellow. Blue lines indicate a 2-fold change and dotted line indicates p>0.05. The number of up or down regulated genes is indicated in blue. The names of the 10 genes with the largest changes are called out. More genes were up-regulated then down regulated in the Δ*yhdP* Δ*fadR* strain and Suppressor 1 vs. wild type. More genes were down regulated in Suppressor 1 compared to the Δ*yhdP* Δ*fadR* strain. **(E)** Relative expression of genes in the CL synthesis pathway is shown as averages ± the SEM and individual data points. No significant changes were evident. **(F)** Relative expression of genes in the fatty acid synthesis and degradation pathways is shown as averages ± the SEM and individual data points. A general normalizing of these pathways occurred in Suppressor 1 compared to the Δ*yhdP* Δ*fadR* strain; however, the most significant change was in the expression of *fabA*. * p<0.05, ** p<0.005, *** p<0.0005 by quasi-linear F-test.

No significant expression changes occurred in the CL biosynthesis pathway (**Fig. 5E**) or other phospholipid biosynthesis genes (**Fig. S9B**). The increased cardiolipin in Δ*yhdP* Δ*fadR* and Suppressor 1 may be post-transcriptional or due to a change in a regulatory pathway not yet altered after 30 minutes at 30 °C (**Fig. S9C**). Levels of mitochondrial CL can be altered due to differences in substrate binding affinity of the CL synthase based on saturation, with increased saturation decreasing CL synthesis (70), and it may be that a similar pathway operates here, albeit in a different direction. We examined the expression of genes in fatty acid synthesis and degradation, many of which are members of the FadR regulon. In Δ*yhdP* Δ*fadR* compared to wild type, many fatty acid biosynthesis genes significantly decreased while many fatty acid degradation genes significantly increased (**Fig. 5F**), consistent with Δ*fadR* effects. While *fabA* expression is decreased, *fabB* expression is not changed, explaining why increased *fabA* expression causes decreased saturation in Suppressor 1. Overall, our data demonstrate the Δ*yhdP* Δ*fadR* strain’s cold sensitivity is due to impaired phospholipid trafficking, and increased phospholipid saturation and cardiolipin are necessary for this functional impairment.

## Discussion

The identification of YhdP, TamB, and YdbH as putative intermembrane phospholipid transporters (30, 31) posed the tantalizing question: why is having three transporters advantageous? Differential phenotypes between the transporters suggest that they possess specialized functions (25, 30, 31, 34, 43-46). Here, we demonstrate each protein plays a distinguishable role in phospholipid transport that can be differentiated based on genetic interactions with lipid metabolism. Disruption of *fadR* and *yhdP* function causes synthetic cold sensitivity not shared by *tamB* or *ydbH*. This sensitivity can be suppressed by removing TamB. This phenotype involves both increased levels of CL and saturated fatty acids, and decreasing amounts of either suppresses. In addition, our data demonstrate that CL synthesized by ClsB and ClsC is qualitatively different—resulting in differential suppression phenotypes—likely due to differences in CL localization, CL-mediated, leaflet-specific changes in lipid packing order (9, 24), localized membrane fluidity (71), or phospholipid transporter interaction.

Based on the currently available data, we cannot rule out the possibility that the suppressors we have identified for the Δ*yhdP* Δ*fadR* cold sensitivity act indirectly (e.g., through regulatory or adaptive changes). However, we now suggest a simple model for the differentiation of function between YhdP and TamB that fully explains our data without invoking additional, more complex interpretations (**Fig. 6A-C**). In this hypothesis, the functions of YhdP and TamB are diversified based on the acyl carbon saturation level preference of the phospholipids they transport: YhdP transports mainly phospholipids with more saturated fatty acids, while TamB transports phospholipids with more unsaturated fatty acids (**Fig. 6A**). Our data and previous data (30, 31) agree YdbH plays a relatively minor role. In the Δ*yhdP* Δ*fadR* strain, YhdP’s absence forces more saturated phospholipids to be transported by TamB. At 30 °C, TamB becomes “clogged” by phospholipids having more saturated fatty acyl side chains, impeding transport (**Fig. 6B**). Bulky and intrinsically disordered CL molecules could contribute to the clogging sterically by obstructing the lipid channel and precluding it from translocating other lipid substrates (i.e., PE). We find it interesting that the phenotype of the Δ*yhdP* Δ*fadR* strain is most severe at low temperature, leading to cold sensitivity. This cold-specific effect may occur due to (i) temperature-dependent decreased membrane fluidity and higher lipid packing order involving lower expression of FabF, which is responsible for increased diunsaturated phospholipids at lower temperature (**Fig. 5F**, **Dataset 3**)(14); (ii) higher relative levels of CL (72); (iii) higher amounts of per cell phospholipids at lower temperatures (72) contributing to increased diffusive flow rate; or, (iv) to thermodynamic properties altering TamB transport specificity and/or substrate behavior contributing to efficiency of substrate loading or transport.

**Figure 6.**
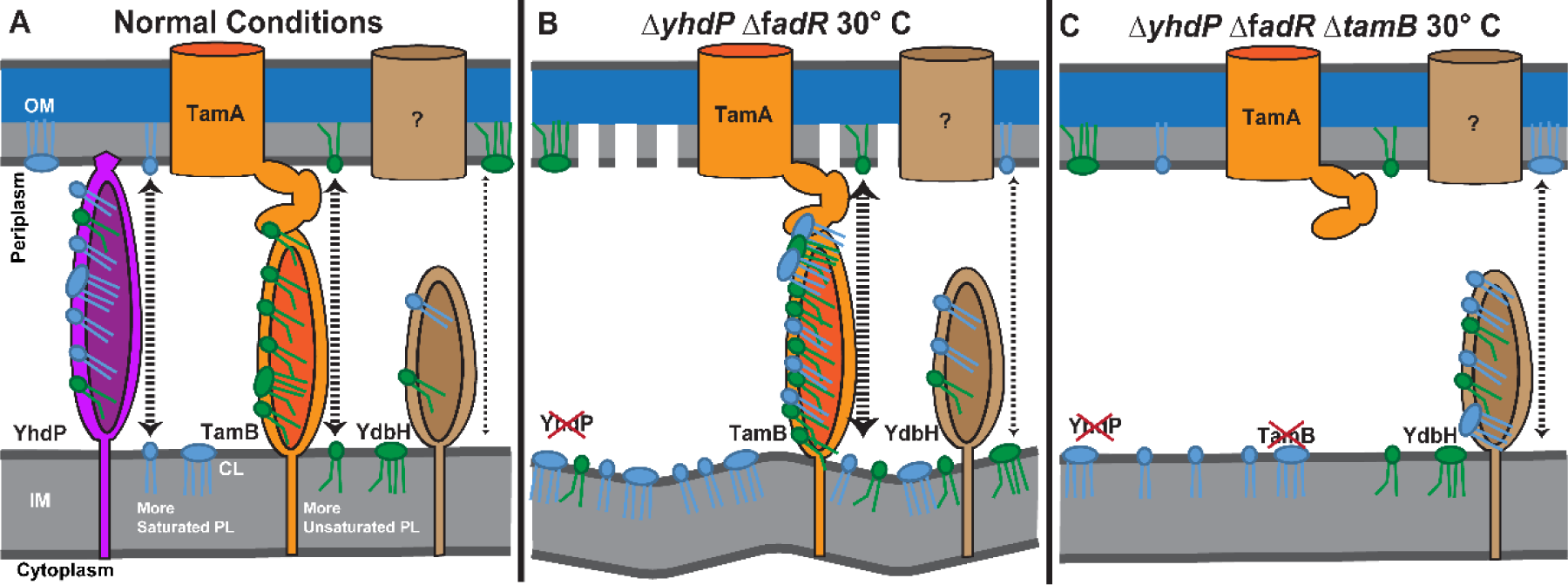
Hypothesized model for the role of YhdP and TamB in phospholipid transport. Our data can be explained by the following model of phospholipid transport. **(A)** In normal conditions, YhdP is mainly responsible for the transport of phospholipids with more saturated fatty acid chains between the IM and OM. TamB shows a strong preference for transporting phospholipids with more unsaturated fatty acids, with YdbH playing a minor role in phospholipid transport. **(B)** In the Δ*yhdP* Δ*fadR* strain, YhdP is not available to transport more saturated phospholipids and the proportion of both saturated fatty acids and cardiolipin is increased, especially at low temperature. The fluidity of the membrane is also lessened by the low temperature. In these conditions, transport of phospholipids between the membranes is impeded, due to clogging of TamB by a combination of less saturated phospholipids and/or bulky and highly disordered CL. This leads to a lethal lack of phospholipids in the OM and overabundance of phospholipids in the IM. **(C)** When *tamB* is deleted in the Δ*yhdP* Δ*fadR* strain, the clogging of TamB is relieved, and the cell relies on YdbH for phospholipid transport overcoming the cold sensitivity. PL: phospholipid

One interesting aspect of our data is that the presence of TamB, with perhaps inhibited function, causes a worse phenotype than the absence of TamB. Furthermore, the phenotype of a Δ*yhdP* Δ*fadR* Δ*ydbH* strain is not measurably worse than that of the Δ*yhdP* Δ*fadR* strain. Together, these data suggest that YdbH is able to successfully transport phospholipids in the Δ*yhdP* Δ*fadR* strain only when TamB is absent. There are several possible mechanisms for this. The first is that YhdP and TamB are the primary transporters and are the preferred interaction partners for other, as of yet unidentified interaction partners (e.g., putative IM proteins facilitating phospholipid transfer to YhdP and TamB). Another possibility is that there is direct toxicity caused by TamB clogging, perhaps through leaking of micellar phospholipids into the periplasm or through stress response overactivation. A third possibility is that YdbH levels are insufficient in the presence of YhdP and TamB to facilitate phospholipid transport and *ydbH* is upregulated in the absence of the other phospholipid transporters. This will be an interesting area for future investigations.

Several pieces of evidence suggest dysregulation of phospholipid transport not just changes to OM phospholipid composition, and so permeability or physical strength, cause the cold sensitivity phenotype of the Δ*yhdP* Δ*fadR* strain. First, the cold sensitivity is unaffected by removing enterobacterial common antigen, which suppresses the membrane permeability of a Δ*yhdP* strain (**Fig. 1C**, (43)), and the OM permeability phenotypes of the Δ*yhdP* Δ*fadR* are not severe (**Fig. S2**). Second, the cold sensitivity is suppressed by deletion of Δ*tamB* (**Fig. 1FG**). Δ*yhdP* Δ*tamB* strains have much greater OM permeability (30) than the Δ*yhdP* or Δ*yhdP* Δ*fadR* strains. Third, CL has been shown to have a normalizing, bidirectional, and leaflet-specific effect on lipid packing order (9, 73-76) and membrane fluidity (77-80) opposite of or similar to that of cholesterol, respectively, and so is unlikely to contribute to a phenotype dependent solely on OM membrane fluidity. Thus, the cold sensitivity phenotype correlates with the impairment of phospholipid trafficking and the likely imbalance of phospholipids between the IM and OM and not with the permeability phenotypes of the OM. Nonetheless, it will be useful in the future to conduction comparative analysis of the phospholipid composition of pure IM and OM extracts (i.e., without cross contamination of the other membrane) at both permissive and non-permissive temperatures to directly link the cold sensitivity observed here with changes in phospholipid transport.

Although both fatty acid saturation and CL levels affect the Δ*yhdP* Δ*fadR* strain’s cold sensitivity, evidence points to saturation rather than headgroup as the main factor separating YhdP and TamB function. Specifically, YhdP loss slows phospholipid transport from the IM to the OM in a *mlaA\** mutant, which loses OM material due to vesiculation and lysis when phospholipid biosynthesis is insufficient to replace lost phospholipids (25, 27). CL only accounts for 5% of phospholipids in exponentially growing cells making it unlikely that the loss of CL transport could slow net phospholipid transport to the extent required to slow lysis.

Our model suggests the IM and OM saturation state could be independently regulated based on YhdP and TamB activity or levels. IM saturation is regulated by altering the composition of newly synthesized fatty acids, while OM saturation could be separately regulated by allowing increased transport of more saturated or unsaturated phospholipids as necessary. IM and OM acyl saturation has been studied in several strains of *E. coli* and in *Yersinia pseudotuberculosis* (7, 19-21) and higher saturated fatty acid levels were observed in the OM than in the IM, supporting discrimination of phospholipid transport based on saturation. The IM needs to maintain an appropriate level of fluidity to allow IM proteins to fold and diffuse, to allow diffusion of hydrophobic molecules, to regulate cellular respiration, and to maintain membrane integrity (84, 85). However, the OM is a permeability barrier, a load-bearing element that resists turgor pressure (1-6), and has an outer leaflet consisting of LPS (2, 8). LPS has saturated acyl chains and is largely immobile in *E. coli* (86) and a specific phospholipid saturation level may be necessary to match or counterbalance LPS rigidity. OM phospholipids may also play a role in the physical strength of the membrane. Interestingly, previous work has demonstrated that the tensile strength of the OM is lessened in a *yhdP* mutant (25).

Possessing two different phospholipid transporters with different saturation state preferences would allow the cell to adapt OM saturation to changing conditions including temperature shifts, chemical and physical insults, or changes in synthesis of other OM components (e.g., LPS or OMPs). A recent study demonstrated in *Y. pseudotuberculosis* the difference in saturation levels between the membranes is exaggerated at 37 °C compared to 8 °C (21). Thus, conditions where IM and OM saturation are independently regulated exist and this regulation is likely important for optimum fitness of the cells. Transcriptional regulation of *yhdP* and *tamB* has not been thoroughly explored. It seems likely *yhdP* transcription is quite complex with several promoters and transcriptional termination sites (87-89). *tamB* is predicted to be in an operon with *tamA* (which codes for an OMP necessary for TamB function) and *ytfP* (87, 90) and to have a σ^E^-dependent promoter (91). The σ^E^ stress response is activated by unfolded OMP accumulation, and downregulates the production of OMPs and upregulates chaperones and folding machinery (74). It is intriguing to speculate a change in OM phospholipid composition might aid in OMP folding due to altered biophysical properties and/or lipoprotein diffusion. In fact, this could explain the observations that some complicated OMPs fold more slowly in the absence of TamB (34, 44-46). Proper phospholipid composition may be necessary for efficient folding of these OMPs, as it is necessary for the correct folding and topology of some IM proteins (73, 92-95). Recently, it has been demonstrated that the *tamAB* operon is regulated by PhoPQ in *Salmonella enterica* Serovar Typhimurium and that this regulation helps maintain OM integrity in the acidic phagosome environment, perhaps due to indirect effects on OMP folding (96). Thus, changing environments may necessitate changes in phospholipid composition that can be accomplished through differential TamB and, perhaps YhdP, regulation.

The mechanism of phospholipid transfer between the IM and OM has been one of the largest remaining questions in gram-negative envelope biogenesis and has remained an area of intense investigation for decades. Furthermore, given the lack of knowledge of this pathway, the role of phospholipid composition in the permeability barrier of the OM, and so the intrinsic antibiotic resistance of gram-negative bacteria, remains unknown. Recent work has begun to elucidate the intermembrane phospholipid transport pathway by identifying YhdP, TamB, and YdbH as proteins capable of transporting phospholipids between the membranes (25, 30, 31). Our work here suggests that YhdP and TamB may have separate roles in phospholipid transport that are differentiated by their preference for phospholipid saturation states and possibly lipid headgroup. Homologs of YhdP and TamB are found throughout *Enterobacterales* and more distantly related members of the AsmA family are widespread in gram-negative bacteria (97). In fact, TamB has recently been shown to play a role in anterograde intermembrane phospholipid transport in the diderm Firmicute *Veillonella parvula* (40, 98) and four AsmA members play a role in phospholipid transport in *Pseudomonas aeruginosa* (41). Structural predictions of YhdP and TamB are strikingly similar between species (32, 33). Given this, it is quite likely similar mechanisms of discrimination in intermembrane phospholipid transfer exist in other species. The data we present here provide a framework for investigation of phospholipid transport and mechanisms differentiating phospholipid transporter function in these species.

## Materials and Methods

### Bacterial strains and Growth Conditions

**Table S3** lists the strains used in this study. Most knockout strains were constructed with Keio collection alleles using P1vir transduction (99, 100). New alleles were constructed through λ-red recombineering (101), using the primers indicated in **Table S4**. The FLP recombinase-FRT system was used to remove antibiotic resistance cassettes as has been described (101). *E. coli* cultures were grown in LB Lennox media at 42°C, unless otherwise mentioned. Where noted, LB was supplemented with vancomycin (Gold Biotechnology), bacitracin (Gold Biotechnology), SDS (Thermo Fisher Scientific), EDTA (Thermo Fisher Scientific), oleic acid (Sigma Aldrich), stearic acid (Sigma Aldrich), arabinose (Gold Biotechnology), or glucose (Thermo Scientific). For plasmid maintenance, media was supplemented with 25 μg/mL kanamycin (Gold Biotechnology), 10 or 20 μg/mL chloramphenicol (Gold Biotechnology), or 10 μg/mL tetracycline (Gold Biotechnology) as appropriate.

### Plasmid construction

The pJW15-P*_fabA_,* and pJW15-P*_fabA_*(-19G>A) reporter plasmids were constructed by amplifying the region upstream of *fabA* from -432 bp to +27 bp relative to the annotated transcriptional start site from wild-type MG1655 or Suppressor 1, respectively, using the Overlap-pJW15-*fabA* FP and RP primers (**Table S4**). pJW15 was amplified using primers Overlap-pJW15-*fabA* FP1 and RP1. The pJW15 plasmid was a kind of gift from Dr. Tracy Raivio (University of Alberta) (102, 103). The resulting fragments were assembled using HiFi Assembly master mix (New England Biolabs) as per the manufacturer’s instructions.

### Growth and Sensitivity Assays

For EOPs assays, cultures were grown overnight, serially diluted, and plated on the indicated media. For *pgsA* depletion, overnight cultures grown in arabinose were washed and diluted 1:100 into fresh LB with 0.2% glucose or arabinose as indicated and grown to OD_600_=1 before diluting and plating on media with glucose or arabinose respective, as above. EOPs were incubated at 30 °C unless otherwise indicated. Growth curves were performed at the indicated temperature as has been described (43), except that starting cultures for growth curves were grown to early stationary phase before inoculation of growth curves rather than overnight. For CPRG assays (50), strains were streaked on LB plates supplemented with CPRG (40µg/ml) and IPTG (100 µM) incubated overnight at 30°C.

### Suppressor isolation and sequencing

Serially diluted cultures were spread on LB plates and incubated overnight at 30°C. Colonies that grew at 30°C were isolated and rechecked for cold resistance through EOP and growth curves. The suppressors were subjected to whole-genome sequencing to identify potential suppressor mutations. Genomic DNA was isolated from the parent and suppressors using a DNeasy Blood and Tissue Kit (Qiagen) as per the manufacturer’s instructions. Library preparation and Illumina sequencing were performed by the Microbial Genome Sequencing Center (MiGS, Pittsburgh, PA, USA). Briefly, Illumina libraries were prepared and the libraries were sequenced with paired-end 151 bp reads on a NextSeq 2000 platform to a read depth of at least 200 Mbp. To identify variants from the reference genome (GenBank: U00096.3), reads were trimmed to remove adaptors and the *breseq* software (version 0.35.4) was used to call variants (104). Identified mutations were confirmed with Sanger sequencing. The consensus motif of the FabR binding site of *fabA* was determined using Multiple Em for Motif Elicitation (MEME) tool in the MEME Suite based on the sequence of putative FabR binding sites reported by Feng and Cronan (2011) (64, 105, 106). The ability of the identified P*_fabA_* mutation to suppress was confirmed by linking the mutant to *zcb*-3059::Tn*10* (107) and transducing to a clean background.

### Lux reporter assays

Overnight cultures of strains harboring the pJW15-P*_fabA_* and pJW15-P*_fabA_*(-19G>A) plasmids were sub-cultured at (1:100) into 200 μl of fresh LB broth in a black 96-well plate. The plate was incubated in a BioTek Synergy H1 plate reader and the OD_600_ and luminescence intensity were recorded every 2.5 min for 6 hours as described in previously (103, 108). Each biological replicate was performed in technical triplicate.

### RNA-seq

Overnight cultures grown at 42°C were subcultured 1:100 into fresh media and incubated at 42 °C until an OD_600_ of 0.4. Then, the cultures were shifted to a 30 °C water bath for 30 minutes. 500 µL of each culture was immediately fixed in the RNAprotect Bacteria Reagent (Qiagen) as per the manufacturer’s instructions. RNA was purified using the RNeasy Kit (Qiagen) with on-column DNase digestion following the manufacturer’s protocol for gram-negative bacteria. Library preparation, Illumina sequencing, and analysis of differentially expressed genes were performed by MiGS. Briefly, samples were treated with RNase free DNase (Invitrogen) and library preparation was performed with the Stranded Total RNA Prep Ligation with Ribo-Zero Plus Kit (Illumina). Libraries were sequenced with 2x50 bp reads on a NextSeq2000 (Illumina). Demultiplexing, quality control, and adapter trimming carried out using bcl-convert (Illumina, v3.9.3) and bcl2fastq (Illumina). Read mapping and read quantification were performed with HISAT2 (109) and the featureCounts tool in Subreader (110), respectively. Descriptive statistics of the RNA-seq read data can be found in **Table S1**. The raw RNA-seq data is available in the Sequence Read Archive database (SRA) (https://www.ncbi.nlm.nih.gov/sra, BioProject ID PRJNA965821; [Reviewer Access](https://dataview.ncbi.nlm.nih.gov/object/PRJNA965821?reviewer=r2dat364brplcph1ant6lu51a3)).

Raw read counts were normalized using the Trimmed Mean of M values algorithm in edgeR (111) and converted to counts per million. The Quasi-Linear F-Test function of edgeR was used for differential expression analysis. Normalized read values and for all genes as well as differential expression analysis statistics can be found in **Dataset S3**. KEGG pathway analysis was conducted using the *“kegga”* function of limma (112). Expression of genes differentially expressed (greater than 2-fold change and p<0.05) was subjected to principle component analysis, and pathway and enrichment analysis using the EcoCyc Omics Dashboard (112).

### Lipid extraction and TLC

For thin layer chromatography (TLC) experiments, cultures were grown in LB with 1 mCi/mL ^32^P until OD_600_ of 0.6-0.8 at the indicated temperature. For temperature downshift experiments, cells were grown at 42 °C until OD_600_ of 0.2 and then transferred to 30 °C for 2 hours. For lipid extraction, cells were pelleted and resuspended in 0.2 mL of 0.5 M NaCl in 0.1 N HCl. Lipids were extracted by first adding 0.6 mL of chloroform:methanol (1:2) to create a single-phase solution. After vigorous vortexing for 30 minutes, 0.2 mL of 0.5 M NaCl in 0.1 N HCl was added to convert the single-phase solution to a two-phase solution. After centrifugation at 20 000 x g for 5 minutes at room temperature, the lower phase was recovered and used for TLC. Approximately 2,000 cpm of phospholipid extract was subjected to TLC analysis using a HPTLC 60 plate (EMD, Gibbstown, NJ)) developed with either solvent 1: chloroform/methanol/acetic acid [60/25/10] (vol/vol/vol) in **Fig. S4A** or by solvent 2: chloroform/methanol/ammonia/water [65/37.5/3/1] (vol/vol/vol/vol) in **Fig. S4B**. Radiolabeled lipids were visualized and quantified using a Phosphoimager Typhoon FLA 9500 (GE). Images were processed and quantified using ImageQuant^TM^ software for scanning and analysis. The phospholipid content is expressed as molecular percentage of the total phospholipids (correcting for the two phosphates per molecule of CL).

### Phospholipid composition analysis

Whole cell lipid extracts were isolated as for TLC temperature downshift experiments without radiolabeling. Absolute amounts of CL, PG, and PE were determined in these lipid extracts using liquid chromatography coupled to electrospray ionization mass spectrometry (LC/MS) in an API 4000 mass spectrometer (Sciex, Framingham, MA, USA) using normal phase solvents as previously published (113). The cardiolipin internal standard (14:0)_4_CL (Avanti Polar Lipids, Alabaster, AL USA) was added to all samples to quantify CL, PG, and PE within a single run. Standard curves containing reference standards of all three phospholipids, (18:1)_4_CL (Avanti Polar Lipids), 16:0_1_18:1_1_PG and 16:0_1_18:1_1_PE (Cayman Chemicals, Ann Arbor, MI, USA) were used to quantify the nmol/mg protein for each species (**Dataset 1**) as previously described (114). Percentage values were calculated by dividing the absolute value of that species by the sum of individual molecular species of CL, PG, or PE. The identities of CL species were confirmed using tandem mass spectrometry.

## Acknowledgments

We thank members of the Mitchell Lab for productive discussions. We would like to thank Professor Tracy Raivio (University of Alberta) for the kind gift of pJW15 plasmid. This work was funded by the National Institute of Allergy and Infectious Disease under Award Number R01-AI155915 and R01-AI155915-S1 (to A.M.M.), the National Institute of General Medical Sciences under Award Number R01-GM121493 (to M.B.), and Texas A&M University start-up funds (to A.M.M.).

## References

1. Nikaido H. Molecular basis of bacterial outer membrane permeability revisited. Microbiology and Molecular Biology Reviews. 2003;67(4):593–656.

2. Silhavy TJ, Kahne D, Walker S. The bacterial cell envelope. Cold Spring Harbor Perspectives in Biology. 2010;2(5).

3. Henderson JC, Zimmerman SM, Crofts AA, Boll JM, Kuhns LG, Herrera CM, Trent MS. The power of asymmetry: Architecture and assembly of the Gram-negative outer membrane lipid bilayer. Annual Review of Microbiology. 2016;70(1):255–78.

4. Berry J, Rajaure M, Pang T, Young R. The spanin complex is essential for lambda lysis. J Bacteriol. 2012;194(20):5667–74.

5. Rojas ER, Billings G, Odermatt PD, Auer GK, Zhu L, Miguel A, et al. The outer membrane is an essential load-bearing element in Gram-negative bacteria. Nature. 2018;559(7715):617-21.

6. Navarro Llorens JM, Tormo A, Martinez-Garcia E. Stationary phase in gram-negative bacteria. FEMS Microbiol Rev. 2010;34(4):476–95.

7. Lugtenberg EJJ, Peters R. Distribution of lipids in cytoplasmic and outer membranes of *Escherichia coli* K12. Biochimica et Biophysica Acta (BBA) - Lipids and Lipid Metabolism. 1976;441(1):38–47.

8. Kamio Y, Nikaido H. Outer membrane of *Salmonella typhimurium*: accessibility of phospholipid head groups to phospholipase c and cyanogen bromide activated dextran in the external medium. Biochemistry. 1976;15(12):2561–70.

9. Bogdanov M, Pyrshev K, Yesylevskyy S, Ryabichko S, Boiko V, Ivanchenko P, et al. Phospholipid distribution in the cytoplasmic membrane of Gram-negative bacteria is highly asymmetric, dynamic, and cell shape-dependent. Science Advances. 2020;6(23):eaaz6333.

10. Benn G, Mikheyeva IV, Inns PG, Forster JC, Ojkic N, Bortolini C, et al. Phase separation in the outer membrane of *Escherichia coli*. Proceedings of the National Academy of Sciences. 2021;118(44):e2112237118.

11. Konovalova A, Silhavy TJ. Outer membrane lipoprotein biogenesis: Lol is not the end. Philosophical Transactions of the Royal Society B: Biological Sciences. 2015;370(1679):20150030.

12. Konovalova A, Kahne DE, Silhavy TJ. Outer Membrane Biogenesis. Annu Rev Microbiol. 2017;71:539–56.

13. Lundstedt E, Kahne D, Ruiz N. Assembly and maintenance of lipids at the bacterial outer membrane. Chemical Reviews. 2021;121(9):5098–123.

14. John E. Cronan J, Rock CO. Biosynthesis of membrane lipids. EcoSal Plus. 2008;3(1).

15. Jones NC, Osborn MJ. Translocation of phospholipids between the outer and inner membranes of *Salmonella typhimurium*. Journal of Biological Chemistry. 1977;252(20):7405–12.

16. Donohue-Rolfe AM, Schaechter M. Translocation of phospholipids from the inner to the outer membrane of *Escherichia coli*. Proceedings of the National Academy of Sciences. 1980;77(4):1867–71.

17. Raetz CR, Kantor GD, Nishijima M, Newman KF. Cardiolipin accumulation in the inner and outer membranes of *Escherichia coli* mutants defective in phosphatidylserine synthetase. J Bacteriol. 1979;139(2):544–51.

18. Langley KE, Hawrot E, Kennedy EP. Membrane assembly: movement of phosphatidylserine between the cytoplasmic and outer membranes of *Escherichia coli*. J Bacteriol. 1982;152(3):1033–41.

19. White DA, Lennarz WJ, Schnaitman CA. Distribution of lipids in the wall and cytoplasmic membrane subfractions of the cell envelope of *Escherichia coli*. Journal of Bacteriology. 1972;109(2):686–90.

20. Ishinaga M, Kanamoto R, Kito M. Distribution of phospholipid molecular species in outer and cytoplasmic membranes of *Escherichia coli*. The Journal of Biochemistry. 1979;86(1):161–5.

21. Davydova L, Bakholdina S, Barkina M, Velansky P, Bogdanov M, Sanina N. Effects of elevated growth temperature and heat shock on the lipid composition of the inner and outer membranes of *Yersinia pseudotuberculosis*. Biochimie. 2016;123:103–9.

22. Osborn MJ, Gander JE, Parisi E, Carson J. Mechanism of Assembly of the Outer Membrane of *Salmonella* typhimurium : Isolation and characterization of cytoplasmic and outer membrane. Journal of Biological Chemistry. 1972;247(12):3962–72.

23. Raetz CR, Newman KF. Diglyceride kinase mutants of *Escherichia coli*: inner membrane association of 1,2-diglyceride and its relation to synthesis of membrane-derived oligosaccharides. J Bacteriol. 1979;137(2):860–8.

24. Bogdanov M. Renovating a double fence with or without notifying the next door and across the street neighbors: why the biogenic cytoplasmic membrane of Gram-negative bacteria display asymmetry?. Emerging Topics in Life Sciences. 2023;7(1):137-50.

25. Grimm J, Shi H, Wang W, Mitchell AM, Wingreen NS, Huang KC, Silhavy TJ. The inner membrane protein YhdP modulates the rate of anterograde phospholipid flow in *Escherichia coli*. Proc Natl Acad Sci U S A 2020;117(43):26907–14.

26. Malinverni JC, Silhavy TJ. An ABC transport system that maintains lipid asymmetry in the Gram-negative outer membrane. Proceedings of the National Academy of Sciences. 2009;106(19):8009–14.

27. Sutterlin HA, Shi H, May KL, Miguel A, Khare S, Huang KC, Silhavy TJ. Disruption of lipid homeostasis in the Gram-negative cell envelope activates a novel cell death pathway. Proceedings of the National Academy of Sciences. 2016;113(11):E1565–E74.

28. May KL, Silhavy TJ. The *Escherichia coli* phospholipase PldA regulates outer membrane homeostasis via lipid signaling. mBio. 2018;9(2):e00379–18.

29. Tang X, Chang S, Qiao W, Luo Q, Chen Y, Jia Z, et al. Structural insights into outer membrane asymmetry maintenance in Gram-negative bacteria by MlaFEDB. Nat Struct Mol Biol. 2021;28(1):81–91.

30. Ruiz N, Davis RM, Kumar S. YhdP, TamB, and YdbH are redundant but essential for growth and lipid homeostasis of the gram-negative outer membrane. mBio. 2021;12(6):e02714–21.

31. Douglass MV, McLean AB, Trent MS. Absence of YhdP, TamB, and YdbH leads to defects in glycerophospholipid transport and cell morphology in Gram-negative bacteria. PLOS Genetics. 2022;18(2):e1010096.

32. Jumper J, Evans R, Pritzel A, Green T, Figurnov M, Ronneberger O, et al. Highly accurate protein structure prediction with AlphaFold. Nature. 2021;596(7873):583-9.

33. Varadi M, Anyango S, Deshpande M, Nair S, Natassia C, Yordanova G, et al. AlphaFold Protein Structure Database: massively expanding the structural coverage of protein-sequence space with high-accuracy models. Nucleic Acids Research. 2021;50(D1):D439–D44.

34. Josts I, Stubenrauch CJ, Vadlamani G, Mosbahi K, Walker D, Lithgow T, Grinter R. The structure of a conserved domain of TamB reveals a hydrophobic β taco fold. Structure. 2017;25(12):1898–906.e5.

35. Kumar N, Leonzino M, Hancock-Cerutti W, Horenkamp FA, Li P, Lees JA, et al. VPS13A and VPS13C are lipid transport proteins differentially localized at ER contact sites. Journal of Cell Biology. 2018;217(10):3625–39.

36. Valverde DP, Yu S, Boggavarapu V, Kumar N, Lees JA, Walz T, et al. ATG2 transports lipids to promote autophagosome biogenesis. J Cell Biol. 2019;218(6):1787–98.

37. Otomo T, Maeda S. ATG2A transfers lipids between membranes *in vitro*. Autophagy. 2019;15(11):2031–2.

38. Osawa T, Kotani T, Kawaoka T, Hirata E, Suzuki K, Nakatogawa H, et al. Atg2 mediates direct lipid transfer between membranes for autophagosome formation. Nat Struct Mol Biol. 2019;26(4):281–8.

39. Cooper BF, Clark R, Kudhail A, Bhabha G, Ekiert DC, Khalid S, Isom GL. Phospholipid transport to the bacterial outer membrane through an envelope-spanning bridge. bioRxiv. 2023:2023.10.05.561070.

40. Grasekamp KP, Beaud Benyahia B, Taib N, Audrain B, Bardiaux B, Rossez Y, et al. The Mla system of diderm Firmicute *Veillonella parvula* reveals an ancestral transenvelope bridge for phospholipid trafficking. Nature Communications. 2023;14(1):7642.

41. Sposato D, Mercolino J, Torrini L, Sperandeo P, Lucidi M, Alegiani R, et al. Redundant essentiality of AsmA-like proteins in *Pseudomonas aeruginosa*. mSphere. 2024;9(2):e00677–23.

42. Mitchell AM, Wang W, Silhavy TJ. Novel Rpos-dependent mechanisms strengthen the envelope permeability barrier during stationary phase. J Bacteriol. 2017;199(2):e00708–16.

43. Mitchell AM, Srikumar T, Silhavy TJ. Cyclic enterobacterial common antigen maintains the outer membrane permeability barrier of *Escherichia coli* in a manner controlled by Yhdp. mBio. 2018;9(4):e01321–18.

44. Selkrig J, Mosbahi K, Webb CT, Belousoff MJ, Perry AJ, Wells TJ, et al. Discovery of an archetypal protein transport system in bacterial outer membranes. Nature Structural & Molecular Biology. 2012;19(5):506–10.

45. Stubenrauch CJ, Dougan G, Lithgow T, Heinz E. Constraints on lateral gene transfer in promoting fimbrial usher protein diversity and function. Open Biology. 2017;7(11):170144.

46. Stubenrauch C, Belousoff MJ, Hay ID, Shen H-H, Lillington J, Tuck KL, et al. Effective assembly of fimbriae in *Escherichia coli* depends on the translocation assembly module nanomachine. Nature Microbiology. 2016;1(7):16064.

47. Cronan JE. The *Escherichia coli* FadR transcription factor: Too much of a good thing? Mol Microbiol. 2021;115(6):1080–5.

48. Nunn WD, Giffin K, Clark D, Cronan JE. Role for *fadR* in unsaturated fatty acid biosynthesis in *Escherichia coli*. Journal of Bacteriology. 1983;154(2):554–60.

49. Rai AK, Mitchell AM. Enterobacterial common antigen: Synthesis and function of an enigmatic molecule. mBio. 2020;11(4):e01914–20.

50. Paradis-Bleau C, Kritikos G, Orlova K, Typas A, Bernhardt TG. A genome-wide screen for bacterial envelope biogenesis mutants identifies a novel factor involved in cell wall precursor metabolism. PLoS Genet. 2014;10(1):e1004056.

51. Ruiz N, Falcone B, Kahne D, Silhavy TJ. Chemical conditionality: a genetic strategy to probe organelle assembly. Cell. 2005;121(2):307–17.

52. Nikaido H, Vaara M. Molecular basis of bacterial outer membrane permeability. Microbiol Rev. 1985;49(1):1–32.

53. Bishop RE. Structural biology of membrane-intrinsic β-barrel enzymes: Sentinels of the bacterial outer membrane. Biochimica et Biophysica Acta (BBA) - Biomembranes. 2008;1778(9):1881–96.

54. Rowlett VW, Mallampalli V, Karlstaedt A, Dowhan W, Taegtmeyer H, Margolin W, Vitrac H. Impact of membrane phospholipid slterations in *Escherichia coli* on cellular function and bacterial stress adaptation. J Bacteriol. 2017;199(13).

55. Tan BK, Bogdanov M, Zhao J, Dowhan W, Raetz CRH, Guan Z. Discovery of a cardiolipin synthase utilizing phosphatidylethanolamine and phosphatidylglycerol as substrates. Proceedings of the National Academy of Sciences. 2012;109(41):16504–9.

56. Pluschke G, Hirota Y, Overath P. Function of phospholipids in *Escherichia coli*. Characterization of a mutant deficient in cardiolipin synthesis. Journal of Biological Chemistry. 1978;253(14):5048–55.

57. Nishijima S, Asami Y, Uetake N, Yamagoe S, Ohta A, Shibuya I. Disruption of the *Escherichia coli cls* gene responsible for cardiolipin synthesis. Journal of Bacteriology. 1988;170(2):775–80.

58. Guo D, Tropp BE. A second *Escherichia coli* protein with CL synthase activity. Biochimica et Biophysica Acta (BBA) - Molecular and Cell Biology of Lipids. 2000;1483(2):263–74.

59. Kikuchi S, Shibuya I, Matsumoto K. Viability of an *Escherichia coli pgsA* null mutant lacking detectable phosphatidylglycerol and cardiolipin. Journal of Bacteriology. 2000;182(2):371–6.

60. Suzuki M, Hara H, Matsumoto K. Envelope disorder of *Escherichia coli* cells lacking phosphatidylglycerol. Journal of Bacteriology. 2002;184(19):5418–25.

61. Shiba Y, Yokoyama Y, Aono Y, Kiuchi T, Kusaka J, Matsumoto K, Hara H. Activation of the Rcs signal transduction system is responsible for the thermosensitive growth defect of an *Escherichia coli* mutant lacking phosphatidylglycerol and cardiolipin. J Bacteriol. 2004;186(19):6526–35.

62. Grabowicz M, Silhavy TJ. Redefining the essential trafficking pathway for outer membrane lipoproteins. Proceedings of the National Academy of Sciences. 2017;114(18):4769–74.

63. Cronan JE, Jr., Gelmann EP. An estimate of the minimum amount of unsaturated fatty acid required for growth of *Escherichia coli*. J Biol Chem. 1973;248(4):1188–95.

64. Feng Y, Cronan JE. Complex binding of the FabR repressor of bacterial unsaturated fatty acid biosynthesis to its cognate promoters. Mol Microbiol. 2011;80(1):195–218.

65. Feng Y, Cronan JE. *Escherichia coli* unsaturated fatty acid synthesis: Complex transcription of the *fabA* gene and *in vivo* identification of the essential reaction catalyzed by FabB. Journal of Biological Chemistry. 2009;284(43):29526–35.

66. Henry MF, Cronan JE. *Escherichia coli* transcription factor that both activates fatty acid synthesis and represses fatty acid degradation. Journal of Molecular Biology. 1991;222(4):843–9.

67. Henry MF, Cronan JE. A new mechanism of transcriptional regulation: Release of an activator triggered by small molecule binding. Cell. 1992;70(4):671–9.

68. Rock CO, Jackowski S. Pathways for the incorporation of exogenous fatty acids into phosphatidylethanolamine in *Escherichia coli*. J Biol Chem. 1985;260(23):12720–4.

69. Heath RJ, Rock CO. Roles of the FabA and FabZ β-hydroxyacyl-acyl carrier protein dehydratases in *Escherichia coli* fatty acid biosynthesis. Journal of Biological Chemistry. 1996;271(44):27795–801.

70. Ostrander DB, Sparagna GC, Amoscato AA, McMillin JB, Dowhan W. Decreased Cardiolipin Synthesis Corresponds with Cytochromec Release in Palmitate-induced Cardiomyocyte Apoptosis. Journal of Biological Chemistry. 2001;276(41):38061–7.

71. Lewis RNAH, Zweytick D, Pabst G, Lohner K, McElhaney RN. Calorimetric, X-Ray Diffraction, and Spectroscopic Studies of the Thermotropic Phase Behavior and Organization of Tetramyristoyl Cardiolipin Membranes. Biophysical Journal. 2007;92(9):3166–77.

72. De Siervo AJ. Alterations in the phospholipid composition of *Escherichia coli* B during growth at different temperatures. J Bacteriol. 1969;100(3):1342–9.

73. Bogdanov M, Sun J, Kaback HR, Dowhan W. A Phospholipid Acts as a Chaperone in Assembly of a Membrane Transport Protein. Journal of Biological Chemistry. 1996;271(20):11615–8.

74. Mitchell AM, Silhavy TJ. Envelope stress responses: balancing damage repair and toxicity. Nature Reviews Microbiology. 2019;17(7):417–28.

75. Luevano-Martinez LA, Kowaltowski AJ. Phosphatidylglycerol-derived phospholipids have a universal, domain-crossing role in stress responses. Arch Biochem Biophys. 2015;585:90–7.

76. Fernandez M, Paulucci NS, Peppino Margutti M, Biasutti AM, Racagni GE, Villasuso AL, et al. Membrane Rigidity and Phosphatidic Acid (PtdOH) Signal: Two Important Events in Acinetobacter guillouiae SFC 500-1A Exposed to Chromium(VI) and Phenol. Lipids. 2019;54(9):557–70.

77. Shibata A, Ikawa K, Shimooka T, Terada H. Significant stabilization of the phosphatidylcholine bilayer structure by incorporation of small amounts of cardiolipin. Biochimica et Biophysica Acta (BBA) - Biomembranes. 1994;1192(1):71–8.

78. Boscia AL, Treece BW, Mohammadyani D, Klein-Seetharaman J, Braun AR, Wassenaar TA, et al. X-ray structure, thermodynamics, elastic properties and MD simulations of cardiolipin/dimyristoylphosphatidylcholine mixed membranes. Chemistry and Physics of Lipids. 2014;178:1–10.

79. Pennington ER, Fix A, Sullivan EM, Brown DA, Kennedy A, Shaikh SR. Distinct membrane properties are differentially influenced by cardiolipin content and acyl chain composition in biomimetic membranes. Biochimica et Biophysica Acta (BBA) - Biomembranes. 2017;1859(2):257–67.

80. Lopes SC, Ivanova G, de Castro B, Gameiro P. Revealing cardiolipins influence in the construction of a significant mitochondrial membrane model. Biochimica et Biophysica Acta (BBA) - Biomembranes. 2018;1860(11):2465–77.

81. Marr AG, Ingraham JL. Effect of temperature on the composition of fatty acids in *Escherichia coli*. Journal of Bacteriology. 1962;84(6):1260–7.

82. Garwin JL, Klages AL, Cronan JE. Beta-ketoacyl-acyl carrier protein synthase II of *Escherichia coli*. Evidence for function in the thermal regulation of fatty acid synthesis. Journal of Biological Chemistry. 1980;255(8):3263–5.

83. de Mendoza D, Klages Ulrich A, Cronan JE. Thermal regulation of membrane fluidity in *Escherichia coli*. Effects of overproduction of beta-ketoacyl-acyl carrier protein synthase I. Journal of Biological Chemistry. 1983;258(4):2098–101.

84. Knapp BD, Huang KC. The effects of temperature on cellular physiology. Annual Review of Biophysics. 2022;51(1):499–526.

85. Bogdanov M, Umeda M, Dowhan W. Phospholipid-assisted refolding of an integral membrane protein: Minimum structural features for phosphatidylethanolamine to act as a molecular chaperone. Journal of Biological Chemistry. 1999;274(18):12339–45.

86. Raetz CR, Whitfield C. Lipopolysaccharide endotoxins. Annu Rev Biochem. 2002;71:635–700.

87. Conway T, Creecy JP, Maddox SM, Grissom JE, Conkle TL, Shadid TM, et al. Unprecedented high-resolution view of bacterial operon architecture revealed by RNA sequencing. mBio. 2014;5(4):e01442–14.

88. Thomason MK, Bischler T, Eisenbart SK, Förstner KU, Zhang A, Herbig A, et al. Global transcriptional start site mapping using differential RNA sequencing reveals novel antisense rnas in *Escherichia coli*. Journal of Bacteriology. 2015;197(1):18–28.

89. Adams PP, Baniulyte G, Esnault C, Chegireddy K, Singh N, Monge M, et al. Regulatory roles of *Escherichia coli* 5’ UTR and ORF-internal RNAs detected by 3’ end mapping. eLife. 2021;10:e62438.

90. Karp PD, Ong WK, Paley S, Billington R, Caspi R, Fulcher C, et al. The EcoCyc database. EcoSal Plus. 2018;8(1):10.1128/ecosalplus.ESP-0006-2018.

91. Huerta AM, Collado-Vides J. Sigma70 promoters in *Escherichia coli*: specific transcription in dense regions of overlapping promoter-like signals. Journal of Molecular Biology. 2003;333(2):261–78.

92. Dowhan W, Bogdanov M. Lipid-dependent membrane protein topogenesis. Annu Rev Biochem. 2009;78:515–40.

93. Bogdanov M, Xie J, Heacock P, Dowhan W. To flip or not to flip: lipid-protein charge interactions are a determinant of final membrane protein topology. J Cell Biol. 2008;182(5):925–35.

94. Bogdanov M, Dowhan W. Lipid-dependent generation of dual topology for a membrane protein. J Biol Chem. 2012;287(45):37939–48.

95. Bogdanov M, Dowhan W, Vitrac H. Lipids and topological rules governing membrane protein assembly. Biochim Biophys Acta. 2014;1843(8):1475–88.

96. Ramezanifard R, Golubeva YA, Palmer AD, Slauch JM. TamAB is regulated by PhoPQ and functions in outer membrane homeostasis during *Salmonella* pathogenesis. J Bacteriol. 2023;205(10):e0018323.

97. Szklarczyk D, Gable AL, Nastou KC, Lyon D, Kirsch R, Pyysalo S, et al. The STRING database in 2021: customizable protein-protein networks, and functional characterization of user-uploaded gene/measurement sets. Nucleic Acids Res. 2021;49(D1):D605–d12.

98. Grasekamp KP, Beaud B, Taib N, Audrain B, Bardiaux B, Rossez Y, et al. Bridges instead of boats? The Mla system of diderm Firmicute Veillonella parvula reveals an ancestral transenvelope core of phospholipid trafficking. bioRxiv. 2023:2023.06.30.547184.

99. Baba T, Ara T, Hasegawa M, Takai Y, Okumura Y, Baba M, et al. Construction of *Escherichia coli* K-12 in-frame, single-gene knockout mutants: the Keio collection. Mol Syst Biol. 2006;2:2006.0008-2006.0008.

100. Silhavy TJ, Berman ML, Enquist LW. Experiments with gene fusions: Cold Spring Harbor Laboratory; 1984.

101. Datsenko KA, Wanner BL. One-step inactivation of chromosomal genes in *Escherichia coli* K-12 using PCR products. Proceedings of the National Academy of Sciences. 2000;97(12):6640–5.

102. MacRitchie DM, Ward JD, Nevesinjac AZ, Raivio TL. Activation of the Cpx envelope stress response down-regulates expression of several locus of enterocyte effacement-encoded genes in enteropathogenic *Escherichia coli*. Infect Immun. 2008;76(4):1465–75.

103. Wong JL, Vogt SL, Raivio TL. Using reporter genes and the *Escherichia coli* ASKA overexpression library in screens for regulators of the Gram negative envelope stress response. Methods Mol Biol. 2013;966:337–57.

104. Deatherage DE, Barrick JE. Identification of mutations in laboratory-evolved microbes from next-generation sequencing data using breseq. Methods Mol Biol. 2014;1151:165–88.

105. Bailey TL, Elkan C. Fitting a mixture model by expectation maximization to discover motifs in biopolymers. Proc Int Conf Intell Syst Mol Biol. 1994;2:28–36.

106. Bailey TL, Johnson J, Grant CE, Noble WS. The MEME Suite. Nucleic Acids Res. 2015;43(W1):W39–49.

107. Singer M, Baker TA, Schnitzler G, Deischel SM, Goel M, Dove W, et al. A collection of strains containing genetically linked alternating antibiotic resistance elements for genetic mapping of Escherichia coli. Microbiol Rev. 1989;53(1):1–24.

108. Rai AK, Carr JF, Bautista DE, Wang W, Mitchell AM. ElyC and cyclic enterobacterial common antigen regulate synthesis of phosphoglyceride-linked enterobacterial common antigen. mBio. 2021;12(6):e02846–21.

109. Kim D, Paggi JM, Park C, Bennett C, Salzberg SL. Graph-based genome alignment and genotyping with HISAT2 and HISAT-genotype. Nature Biotechnology. 2019;37(8):907–15.

110. Liao Y, Smyth GK, Shi W. featureCounts: an efficient general purpose program for assigning sequence reads to genomic features. Bioinformatics. 2013;30(7):923–30.

111. Robinson MD, McCarthy DJ, Smyth GK. edgeR: a Bioconductor package for differential expression analysis of digital gene expression data. Bioinformatics. 2009;26(1):139–40.

112. Ritchie ME, Phipson B, Wu D, Hu Y, Law CW, Shi W, Smyth GK. limma powers differential expression analyses for RNA-sequencing and microarray studies. Nucleic Acids Research. 2015;43(7):e47-e.

113. Sparagna GC, Johnson CA, McCune SA, Moore RL, Murphy RC. Quantitation of cardiolipin molecular species in spontaneously hypertensive heart failure rats using electrospray ionization mass spectrometry. J Lipid Res. 2005;46(6):1196–204.

114. Hryc CF, Mallampalli VKPS, Bovshik EI, Azinas S, Fan G, Serysheva II, et al. Structural insights into cardiolipin replacement by phosphatidylglycerol in a cardiolipin-lacking yeast respiratory supercomplex. Nature Communications. 2023;14(1):2783.

