## Supplemental Material for "Genetic evidence for functional diversification of gram-negative intermembrane phospholipid transporters"

**This file includes:**

Figures S1 to S9

Tables S1 to S4

Legends for Datasets S1 to S3

Supplemental References

**Other supplementary materials for this manuscript:**

Datasets S1 to S4

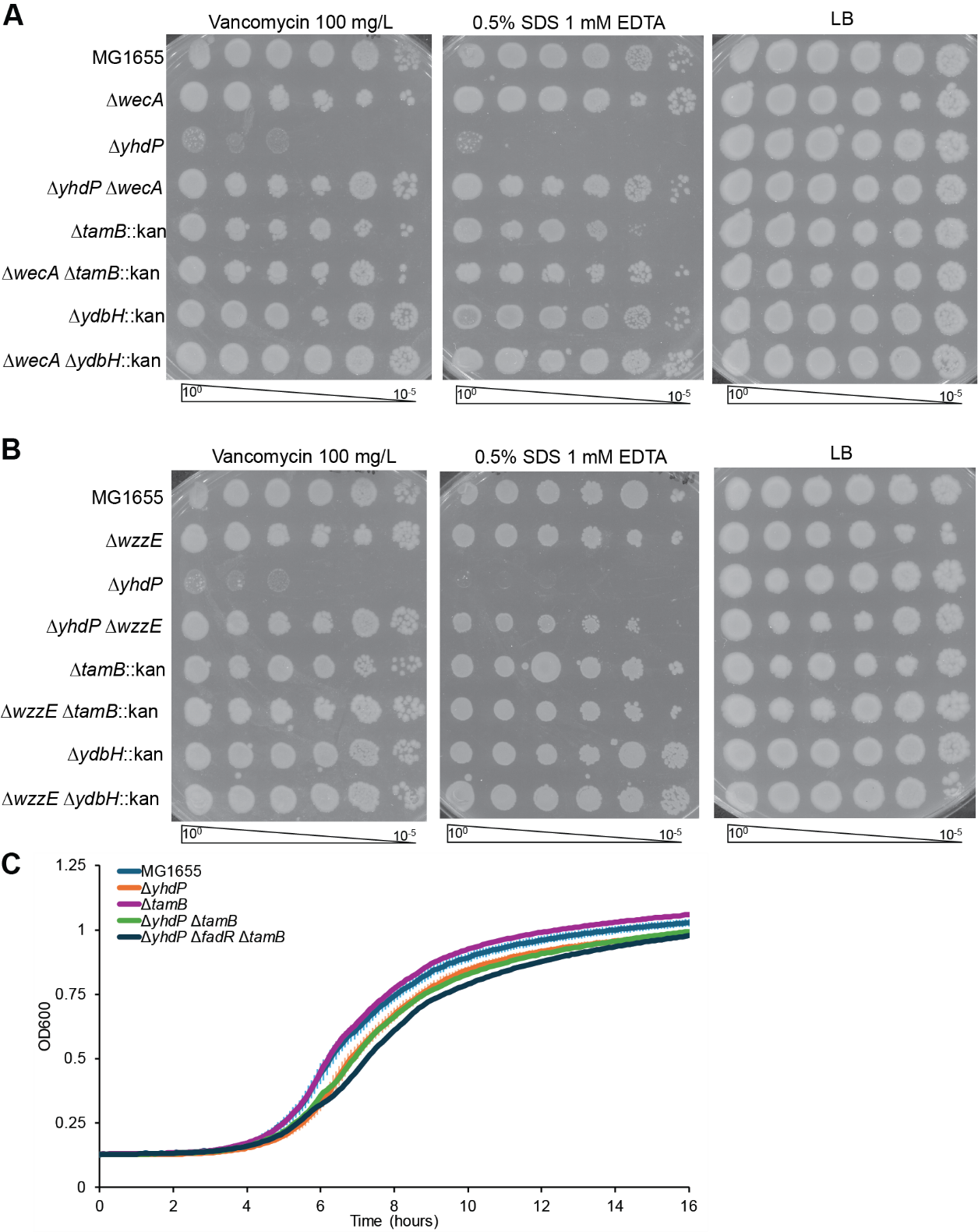

**Fig S1. Effects of *yhdP,* *tamB,* and *ydbH* on outer membrane permeability and growth.** **(A-B)** Efficiency of plating assays (EOPs) were performed at 37 °C in the indicated conditions. Only ∆*yhdP* causes significant outer membrane (OM) permeability to vancomycin and SDS (sodium dodecyl sulfate) EDTA. Loss of all enterobacterial common antigen due to ∆*wecA* deletion (**A**) or loss of cyclic enterobacterial common antigen due to ∆*wzzE* **(B)** suppresses ∆*yhdP* OM permeability but does not change ∆*tamB* or ∆*ydbH* phenotypes. Data are representative of three independent experiments.

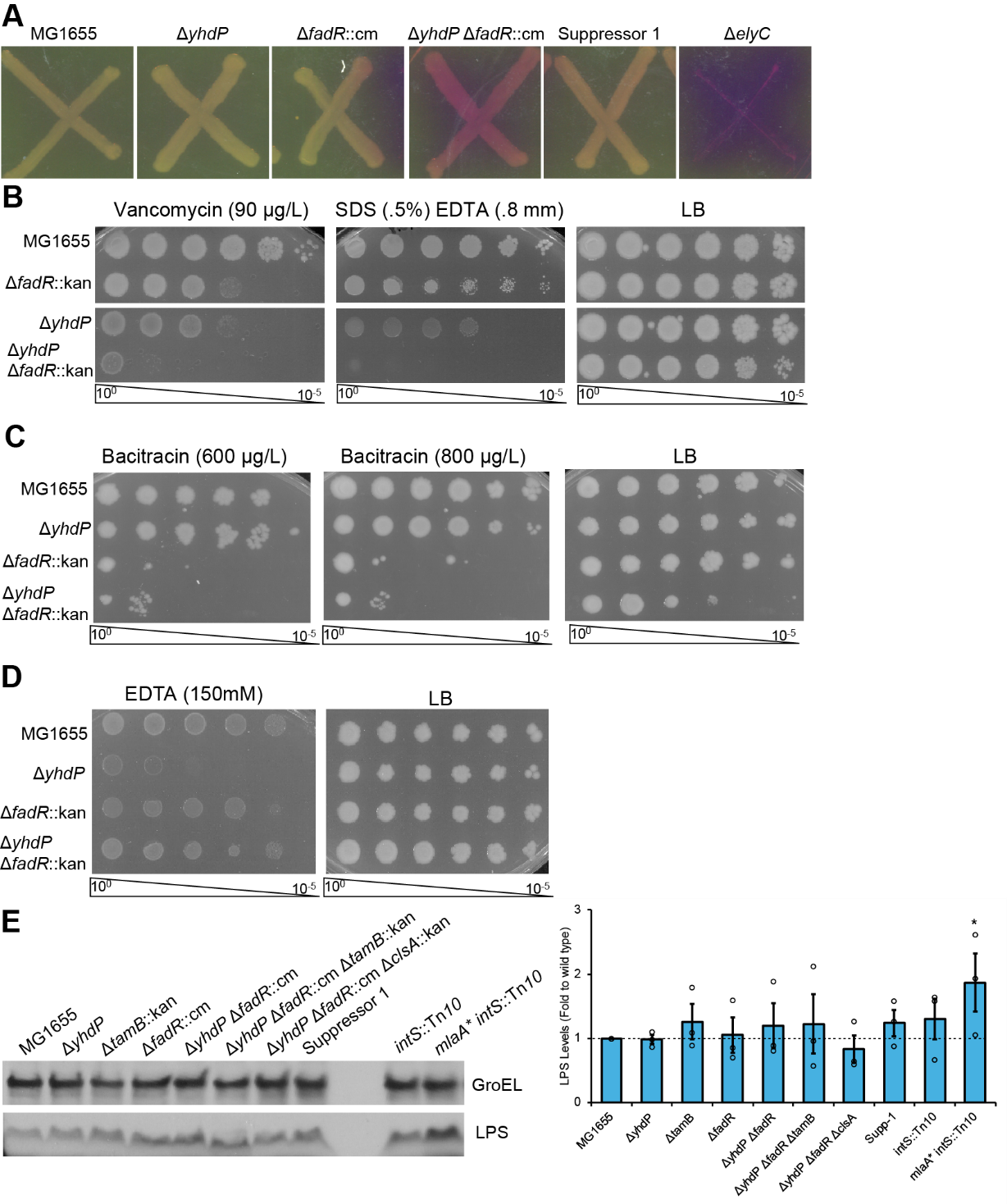
**Figure S2. Envelope permeability phenotypes of ∆*yhdP* ∆*fadR* mutants.** **(A)** A CPRG assay was used to assay lysis and envelope permeability. Red color indicates the production of chlorophenol red after β-galactosidase cleavage of CPRG. The ∆*yhdP* ∆*fadR* strain shows increased lysis or envelope permeability compared to the wild type strain and single mutants. The ∆*elyC* strain serves as a positive control. **(B-D)** Sensitivity of strains to various compounds inhibited by the OM was assayed by EOP at 37 °C. **(B)** The ∆*yhdP* ∆*fadR* strain shows additive sensitivity to vancomycin and SDS EDTA when compared to its parent strains. **(C)** The ∆*yhdP* ∆*fadR* strain demonstrates similar bacitracin sensitivity to the ∆*fadR* strain. **(D)** The ∆*yhdP* ∆*fadR* strain does not exhibit the EDTA sensitivity of a ∆*yhdP* strain. **(E)** LPS levels in cultures grown to OD=0.2 at 37 °C then down shifted to 30 °C for two hours were assayed by immunoblot analysis (α-LPS core, Hycult Biotechnology) as has been described (1-3). GroEL (Millipore Sigma) serves as a loading control. The *mlaA** strain has increased LPS levels and serves as a control. Relative LPS levels from three biological replicates were determined by densitometry and are shown as the mean ± the SEM with individual data points. The *mlaA** strain has the only significant change in LPS levels. All images are representative of three independent experiments.

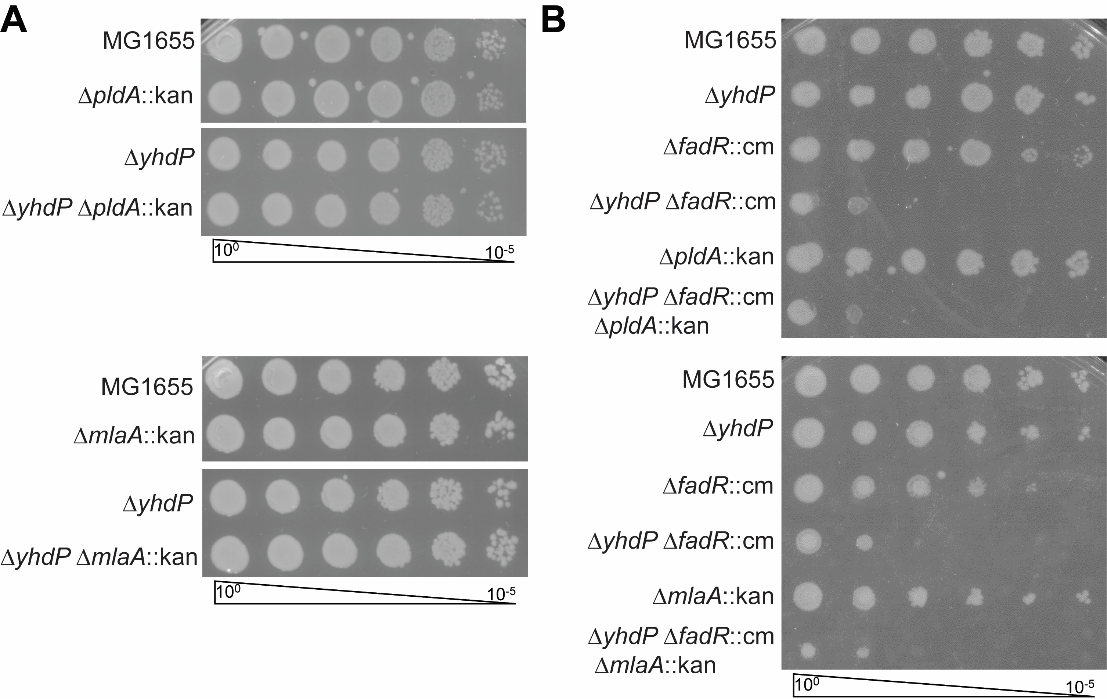

**Fig. S3. *yhdP* does not show genetic interaction with OM asymmetry mutants.** EOPs were carried out at 30 °C on LB media. **(A)** Combination of ∆*yhdP* with ∆*pldA* or ∆*mlaA* did not result in cold sensitivity. **(B)** Combination of ∆*yhdP* ∆*fadR* with ∆*pldA* or ∆*mlaA* did not suppress cold sensitivity. Images are representative of three independent experiments.

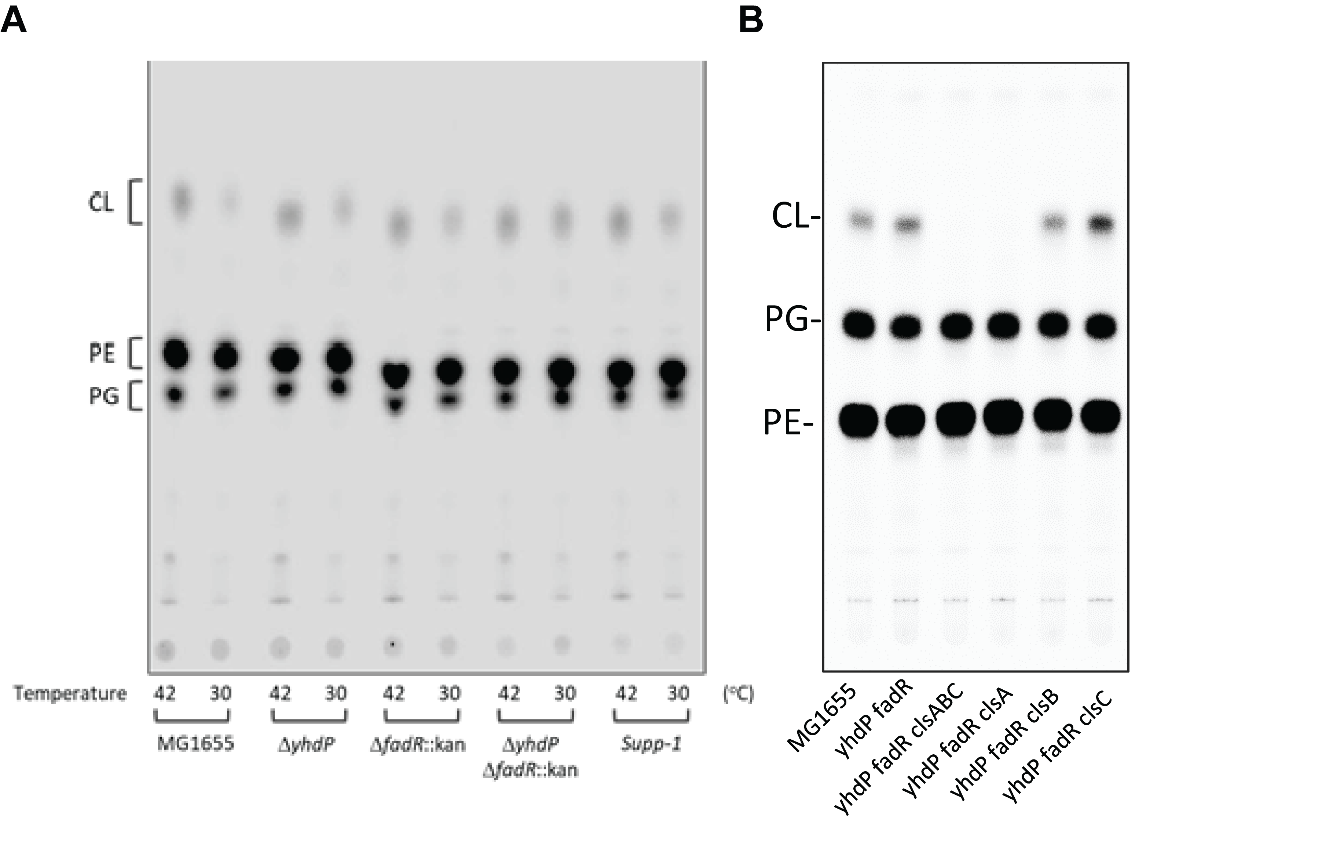

**Fig. S4. Representative TLC images.** Cultures of the indicated strains were grown to log phase at 42 °C then transferred to 30 °C or 42 °C for 2 hours before performing thin layer chromatography to analyze phospholipid content. Two representative images are shown. Solvent conditions are different between image (A) (chloroform-methanol-acetic acid [60/25/10] (vol/vol/vol)) and (B) (chloroform-methanol-ammonia-water [65/37.5/3/1] (vol/vol/vol/vol)) to allow better separation of PG and PE. CL: cardiolipin; PE: phosphatidylethanolamine; PG: phosphatidylglycerol.

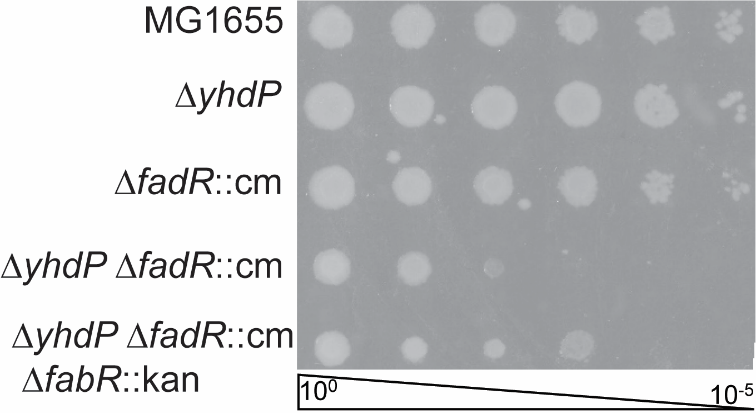

**Fig. S5. Deletion of *fabR* partially suppresses the cold sensitivity of the ∆*yhdP* ∆*fadR* strain.** EOPs were performed at 30 °C on LB media with the indicated strains. Deletion of *fabR* partially suppresses the cold sensitivity of the ∆*yhdP* ∆*fadR* strain. Image is representative of three independent experiments.

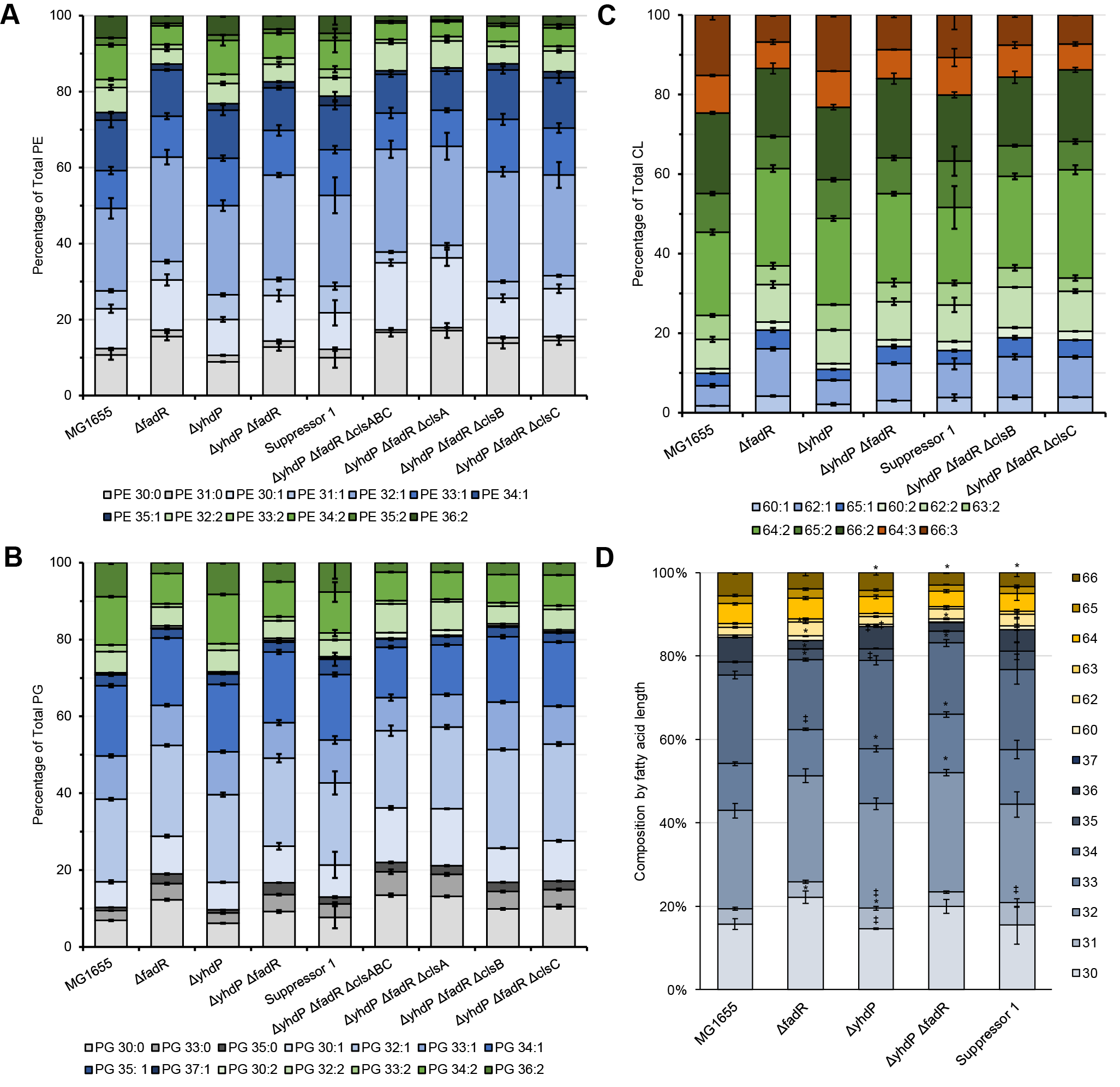

**Fig. S6. Full phospholipid composition for down shifted strains.** Phospholipid composition indicted strains was assayed using LC/MS after growth to OD_600­_=0.2 at 42 °C then down shifting the temperature to 30 °C for 2 hours. **(A-C)** Percentages of PE (A), PG (B), and CL (C) were calculated from absolute quantification of all species detected. Data for phospholipids without unsaturations are shown in grey, with one unsaturation in blue, with two unsaturations in green, with three unsaturations in orange. **(D)** Total fatty acid lengths for all phospholipids are shown with CL specific lengths shown in yellow tones. * p<0.05 vs. MG1655 by the Mann-Whitney test; ‡ p<0.05 vs. the ∆*yhdP* ∆*fadR* strain by the Mann-Whitney test.

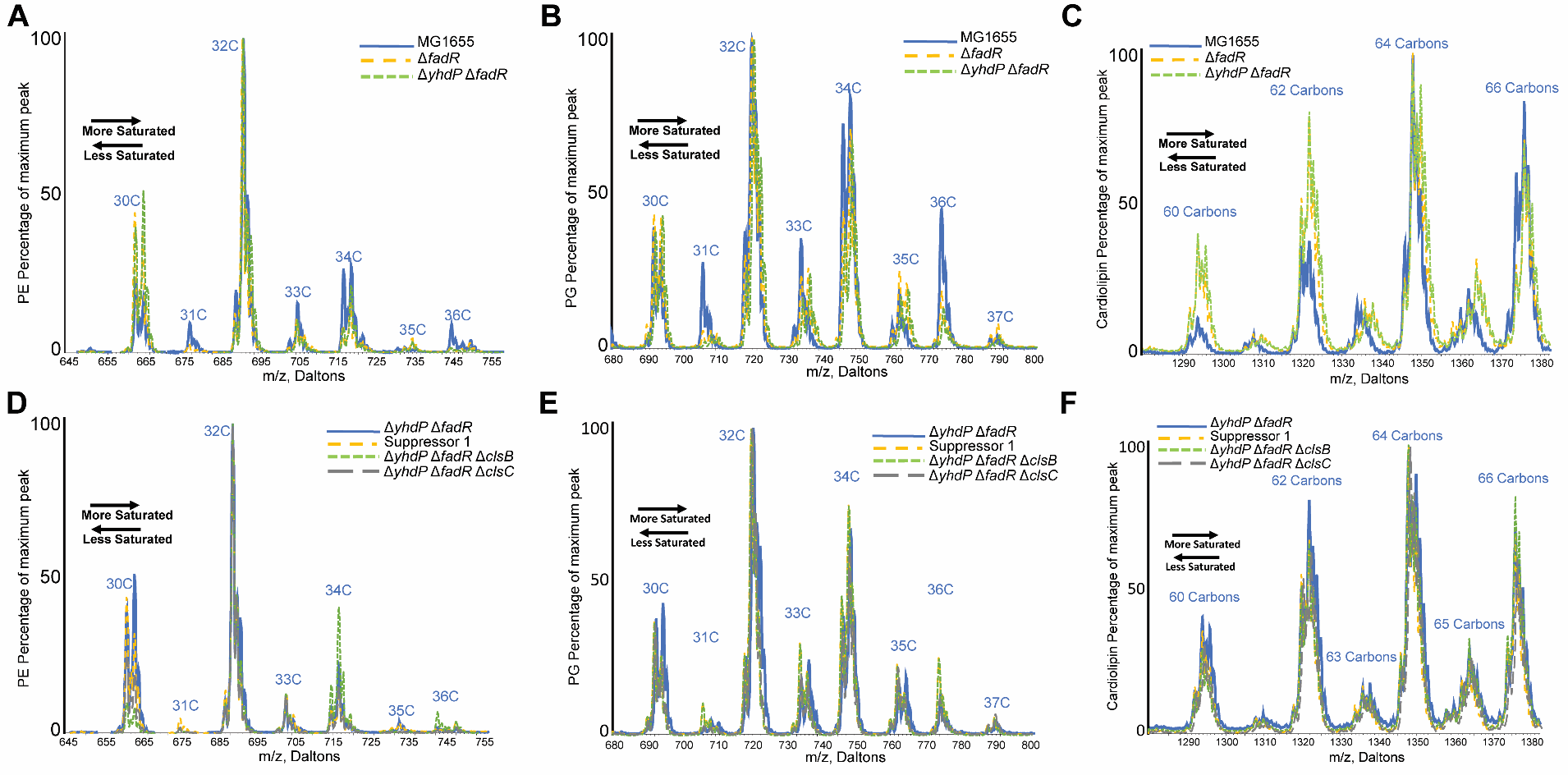

**Fig. S7. Example MS spectra for phospholipid composition.** Phospholipid composition indicted strains was assayed using LC/MS. Representative spectra are shown the wild type, Δ*fadR*, and Δ*yhdP* Δ*fadR* strains (A-C) and for the Δ*yhdP* Δ*fadR*, Suppressor 1, Δ*yhdP* Δ*fadR* Δ*clsB*, and Δ*yhdP* Δ*fadR* Δ*clsC* strains (D-F). Spectra for PE (A, D), PG (B, E), and CL (C, F) are shown as separate panels. Data are shown as relative values to the maximum peak height.

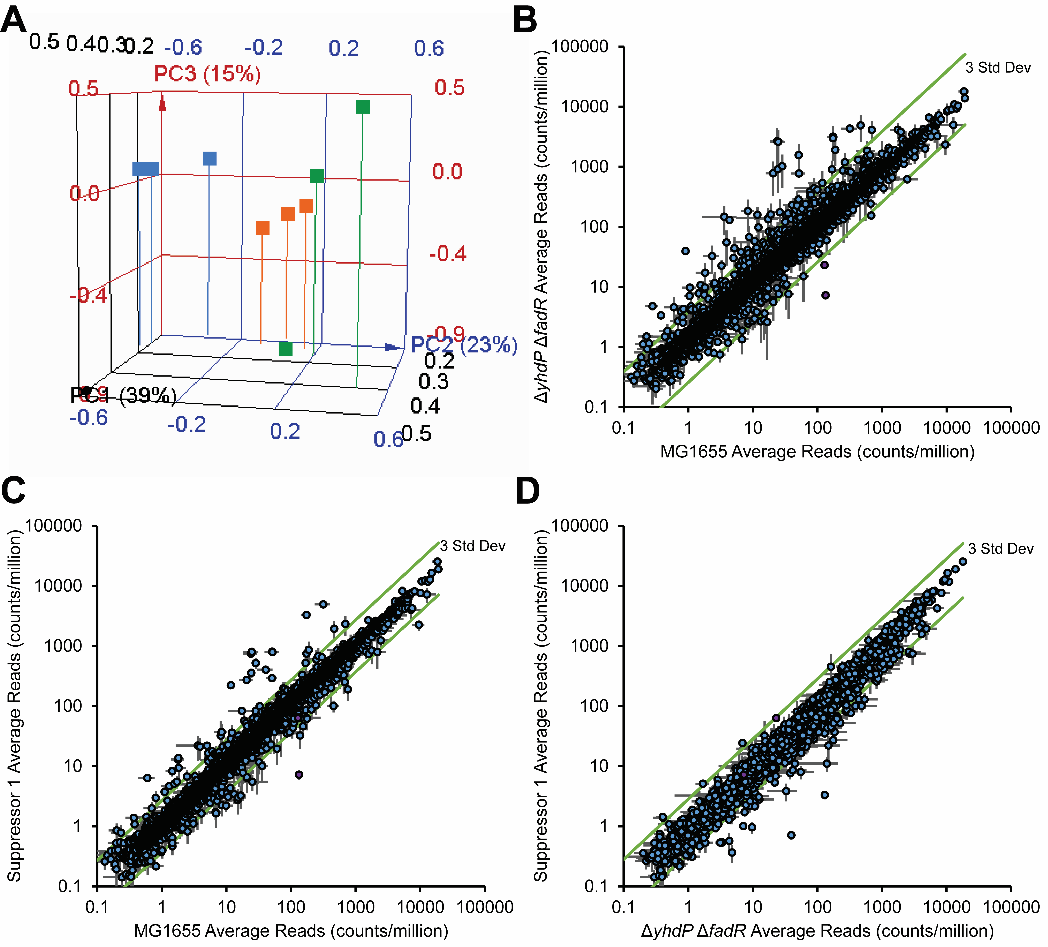

**Fig. S8. RNA-seq read data.** RNA-seq was performed for the wild-type, ∆*yhdP* ∆*fadR*, and Suppressor 1 strains (see main text for details). **(A)** Principal component analysis was performed for the expression of all genes differentially expressed between any two groups. The samples are graphed in relation to the top three principle components. Blue: wild type; Green: ∆*yhdP* ∆*fadR*; Orange: Suppressor 1. Suppressor 1 and the ∆*yhdP* ∆*fadR* strain cluster most closely together. **(B-D)** The mean read count per gene ± the SEM is shown for **(A)** the ∆*yhdP* ∆*fadR* strain vs. wild type, **(C)** Suppressor 1 vs. wild type, and **(D)** Suppressor 1 vs. the ∆*yhdP* ∆*fadR* strain.

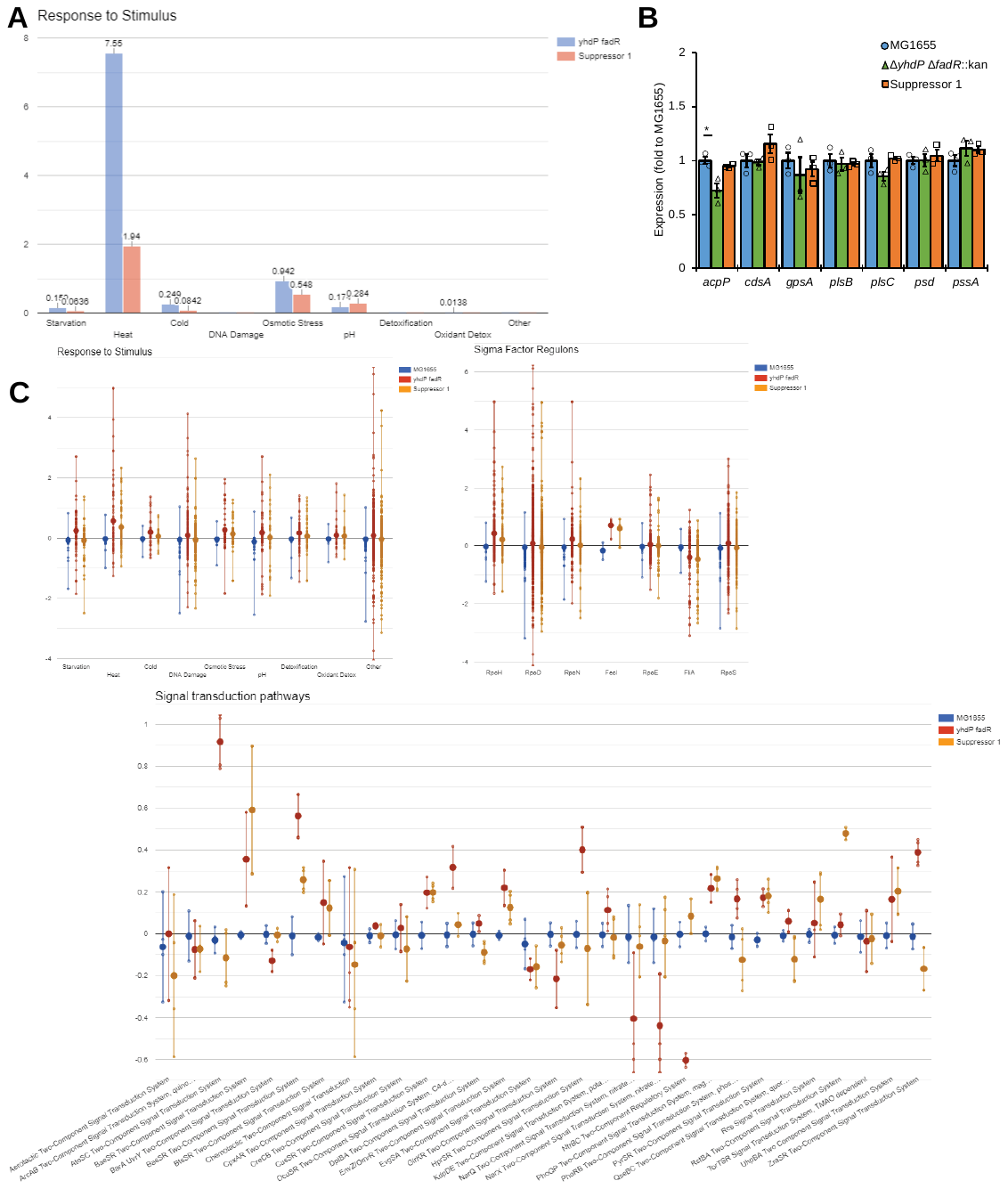

**Fig. S9. RNA-seq enrichment and pathway analysis. (A)** Pathway enrichment analysis was performed on RNA-seq data from the ∆*yhdP* ∆*fadR* strain and Suppressor 1 against wild type using the EcoCyc Omics Dashboard. Data for genes in responses to external stimuli are shown. Graphed values at –log(p-values) calculated by a Fisher-exact test. **(B)** Relative expression of genes in phospholipid biosynthesis not specific for CL synthesis. Data are shown as mean ± SEM and individual data points. * p<0.05 by quasi-linear F-test. **(C)** Pathway analysis was performed using the EcoCyc Omics Dashboard on the expression of all genes. Data for response to stimulus, sigma factor regulons, and signal transduction pathways are shown as fold values. Large dots indicate the mean for the pathway while small dots indicate individual genes. Lines indicate the range of changes for the pathway.

**Table S1: RNA-Seq Descriptive Statistics**

| **Read Alignment Statistics** | | | | | |
| --- | --- | --- | --- | --- | --- |
| **Sample** | **Total Reads** | **All Mapped Reads** | **Coding sequence Mapped Reads** | **rRNA Mapped Reads** | **tRNA Mapped Reads** |
| **MG1655 1** | 35,023,454 | 35,000,224 | 29,647,965 | 27,966 | 92,803 |
| **MG1655 2** | 34,950,112 | 34,920,774 | 28,513,228 | 28,134 | 245,155 |
| **MG1655 3** | 36,311,924 | 36,277,404 | 30,568,531 | 43,327 | 180,315 |
| **∆*yhdP* ∆*fadR* 1** | 36,218,546 | 36,052,383 | 30,202,733 | 28,331 | 164,775 |
| **∆*yhdP* ∆*fadR* 2** | 35,056,248 | 34,914,159 | 27,808,209 | 24,290 | 251,332 |
| **∆*yhdP* ∆*fadR* 3** | 34,920,436 | 34,789,325 | 28,934,121 | 36,209 | 189,229 |
| **Suppressor 1 1** | 33,074,066 | 32,998,982 | 27,683,364 | 27,599 | 120,694 |
| **Suppressor 1 2** | 32,258,252 | 32,189,270 | 26,838,003 | 23,141 | 180,518 |
| **Suppressor 1 3** | 33,379,798 | 33,330,014 | 21,515,004 | 612,947 | 108,837 |
| **Median** | **34,950,112** | **34,914,159** | **28,513,228** | **28,134** | **180,315** |

| **Log2 Fold Values Between Indicated Sample Groups** | | | |
| --- | --- | --- | --- |
| **Statistic** | **∆*yhdP* ∆*fadR*/ MG1655** | **Suppressor 1/ ∆*yhdP* ∆*fadR*** | **Suppressor 1/ MG1655** |
| **Mean** | 0.139 | -0.134 | 0.005 |
| **Median** | 0.055 | -0.052 | -0.012 |
| **Standard Deviation** | 0.650 | 0.499 | 0.477 |
| **Maximum** | 6.80 | 3.61 | 5.02 |
| **Minimum** | -4.20 | -5.78 | -4.21 |

**Table S2: RNA-Seq KEGG Pathway Enrichment Analysis**

| **Pathway Identifier** | **Pathway** | **N** | **MG1655 vs ∆*yhdP ∆fadR*** | | | | **∆*yhdP ∆fadR* vs. Suppressor 1** | | | | **MG1655 vs. Suppressor 1** | | | |
| --- | --- | --- | --- | --- | --- | --- | --- | --- | --- | --- | --- | --- | --- | --- |
|  |  |  | **Up** | **Down** | **P. Up** | **P. Down** | **Up** | **Down** | **P. Up** | **P. Down** | **Up** | **Down** | **P. Up** | **P. Down** |
| **path:eco00020** | **Citrate cycle (TCA cycle)** | 29 | 4 | 0 | 0.003 | 1 | 0 | 0 | 1 | 1 | 0 | 0 | 1 | 1 |
| **path:eco00190** | **Oxidative phosphorylation** | 43 | 4 | 0 | 0.012 | 1 | 0 | 0 | 1 | 1 | 0 | 0 | 1 | 1 |
| **path:eco00410** | **beta-Alanine metabolism** | 14 | 2 | 0 | 0.033 | 1 | 0 | 0 | 1 | 1 | 1 | 0 | 0.05571 | 1 |
| **path:eco00620** | **Pyruvate metabolism** | 59 | 5 | 0 | 0.007 | 1 | 0 | 0 | 1 | 1 | 1 | 0 | 0.215643 | 1 |
| **path:eco00650** | **Butanoate metabolism** | 35 | 7 | 0 | 5.68E-06 | 1 | 0 | 1 | 1 | 0.148376 | 1 | 0 | 0.133825 | 1 |
| **path:eco01220** | **Degradation of aromatic compounds** | 17 | 2 | 0 | 0.048 | 1 | 0 | 1 | 1 | 0.074888 | 0 | 0 | 1 | 1 |
| **path:eco02026** | **Biofilm formation - Escherichia coli** | 52 | 4 | 0 | 0.022 | 1 | 0 | 0 | 1 | 1 | 0 | 0 | 1 | 1 |
| **path:eco00627** | **Aminobenzoate degradation** | 6 | 1 | 0 | 0.118978 | 1 | 0 | 1 | 1 | 0.027 | 0 | 0 | 1 | 1 |
| **path:eco00630** | **Glyoxylate and dicarboxylate metabolism** | 42 | 2 | 0 | 0.218309 | 1 | 0 | 0 | 1 | 1 | 2 | 0 | 0.012 | 1 |
| **path:eco01212** | **Fatty acid metabolism** | 21 | 5 | 1 | 5.53E-05 | 0.073103 | 0 | 0 | 1 | 1 | 3 | 1 | 7.17E-05 | 0.020 |
| **path:eco00061** | **Fatty acid biosynthesis** | 13 | 0 | 1 | 1 | 0.046 | 0 | 0 | 1 | 1 | 0 | 1 | 1 | 0.012 |
| **path:eco00071** | **Fatty acid degradation** | 15 | 5 | 0 | 9.00E-06 | 1 | 0 | 0 | 1 | 1 | 3 | 0 | 2.49E-05 | 1 |
| **path:eco00280** | **Valine, leucine and isoleucine degradation** | 11 | 4 | 0 | 5.23E-05 | 1 | 0 | 0 | 1 | 1 | 2 | 0 | 8.43E-04 | 1 |
| **path:eco00281** | **Geraniol degradation** | 6 | 4 | 0 | 2.58E-06 | 1 | 0 | 0 | 1 | 1 | 2 | 0 | 2.33E-04 | 1 |
| **path:eco00362** | **Benzoate degradation** | 12 | 5 | 0 | 2.50E-06 | 1 | 0 | 0 | 1 | 1 | 2 | 0 | 0.001 | 1 |
| **path:eco00380** | **Tryptophan metabolism** | 10 | 2 | 0 | 0.017 | 1 | 0 | 0 | 1 | 1 | 1 | 0 | 0.040 | 1 |
| **path:eco00592** | **alpha-Linolenic acid metabolism** | 3 | 2 | 0 | 0.001 | 1 | 0 | 0 | 1 | 1 | 1 | 0 | 0.012 | 1 |
| **path:eco00903** | **Limonene and pinene degradation** | 3 | 2 | 0 | 0.001 | 1 | 0 | 0 | 1 | 1 | 1 | 0 | 0.012 | 1 |
| **path:eco00930** | **Caprolactam degradation** | 3 | 2 | 0 | 0.001 | 1 | 0 | 0 | 1 | 1 | 1 | 0 | 0.012 | 1 |
| **path:eco01110** | **Biosynthesis of secondary metabolites** | 338 | 12 | 0 | 0.047 | 1 | 0 | 0 | 1 | 1 | 4 | 0 | 0.043 | 1 |
| **path:eco01120** | **Microbial metabolism in diverse environments** | 268 | 14 | 0 | 0.001 | 1 | 0 | 2 | 1 | 0.348026 | 4 | 0 | 0.020 | 1 |
| **path:eco01200** | **Carbon metabolism** | 110 | 8 | 0 | 0.002 | 1 | 0 | 0 | 1 | 1 | 3 | 0 | 0.009 | 1 |

**Table S3: Strains used in this study**

| **Strain** | **Genotyping** | **Reference** |
| --- | --- | --- |
| MG1655 | K-12 F^-^ λ^-^ *rph-1* | (4) |
| AM182 | MG1655 Δ*yhdP* | (2) |
| AK38 | MG1655 Δ*fadR*::cm | This study |
| AK31 | MG1655 Δ*fadR*::kan | This study |
| AK7 | MG1655 Δ*yhdp* Δ*fadR*::kan | This study |
| AK58 | MG1655 Δ*yhdp* Δ*fadR*::cm | This study |
| AM174 | MG1655 Δ*wecA*::kan | (5) |
| AM334 | MG1655 Δ*wecA* | (2) |
| AM365 | MG1655 Δ*wzzE* | (2) |
| AK183 | MG1655 Δ*yhdp* Δ*fadR*::cm Δ*wecA*::kan | This study |
| AM1442 | MG1655 Δ*tamB*::kan | This study |
| AK176 | MG1655 Δ*ydbH*::kan | This study |
| AM341 | MG1655 Δ*yhdP* Δ*wecA* | (2) |
| AM369 | MG1655 Δ*yhdP* Δ*wzzE* | (2) |
| AM1444 | MG1655 Δ*wecA* Δ*tamB*::kan | This study |
| AM1445 | MG1655 Δ*wecA* Δ*ydbH*::kan | This study |
| AM1446 | MG1655 Δ*wzzE* Δ*tamB*::kan | This study |
| AM1447 | MG1655 Δ*wzzE* Δ*ydbH*::kan | This study |
| AK113 | MG1655 Δ*tamB*::kan Δ*fadR*::cm | This study |
| AK177 | MG1655 Δ*ydbH*::kan Δ*fadR*::cm | This study |
| AM743 | MG1655 Δ*elyC* | (3) |
| AK3 | MG1655 Δ*yhdp* Δ*fadR*::kan P*_fabA_*(-19G>A) (Suppressor 1) | This study |
| AK200 | MG1655 Δ*yhdp* Δ*fadR*::kan *zcb*-3059::Tn*10* | This study; Tn*10* allele from (6) |
| AK201 | MG1655 Δ*yhdp* Δ*fadR*::kan *zcb*-3059::Tn*10* P*_fabA_*(-19G>A) | This study |
| AM384 | MG1655 Δ*pldA*::kan | This study |
| AM385 | MG1655 Δ*yhdp* Δ*pldA*::kan | This study |
| AM522 | MG1655 Δ*mlaA*::kan | (2) |
| AM524 | MG1655 Δ*yhdp* Δ*mlaA*::kan | This study |
| AK182 | MG1655 Δ*yhdp* Δ*fadR*::cm Δ*pldA*::kan | This study |
| AK180 | MG1655 Δ*yhdp* Δ*fadR*::cm Δ*mla*::kan | This study |
| AK167 | MG1655 Δ*clsA*::kan | This study |
| AK168 | MG1655 Δ*clsB*::kan | This study |
| AK169 | MG1655 Δ*clsC*::kan | This study |
| AM1357 | MG1655 Δ*clsA* Δ*clsB* Δ*clsC* | This study |
| AK171 | MG1655 Δ*yhdP* Δ*fadR*::cm Δ*clsA*::kan | This study |
| AK172 | MG1655 Δ*yhdP* Δ*fadR*::cm Δ*clsB*::kan | This study |
| AK173 | MG1655 Δ*yhdP* Δ*fadR*::cm Δ*clsC*::kan | This study |
| AK274 | MG1655 Δ*clsA* Δ*clsB* Δ*clsC* Δ*yhdP* Δ*fadR*::cm | This study |
| AM1409 | MG1655 Δ*rcsF* Δ*lpp* Δ*pgsA* pBAD33-pgsA | (7) |
| AK205 | MG1655 Δ*rcsF* Δ*lpp* Δ*pgsA* Δ*yhdP*::kan *fadR*::Tn*10* pBAD33-pgsA | This study |
| AK89 | MG1655 Δ*fabR*::kan | This study |
| AK84 | MG1655 Δ*yhdP* Δ*fadR*::cm Δ *fabR*::kan | This study |
| AK178 | MG1655 Δ*yhdP* Δ*fadR*::cm Δ*tamB*::kan | This study |
| AM1469 | MG1655 Δ*yhdP* Δ*tamB*::kan | This study |
| AK179 | MG1655 Δ*yhdP* Δ*fadR*::cm Δ*ydbH*::kan | This study |
| AM944 | MG1655 pJW15 | (3); plasmid from (8) |
| AK140 | MG1655 pJW15-P*_fabA_* | This study |
| AK141 | MG1655 pJW15-P*_fabA_*(-19G>A) | This study |
| AK142 | MG1655 Δ*yhdP* pJW15-P*_fabA_* | This study |
| AK143 | MG1655 Δ*yhdP* pJW15-P*_fabA_*(-19G>A) | This study |
| AK144 | MG1655 Δ*fadR*::cm pJW15-P*_fabA_* | This study |
| AK145 | MG1655 Δ*fadR*::cm pJW15-P*_fabA_*(-19G>A) | This study |
| AK146 | MG1655 Δ*yhdP* Δ*fadR*::cm pJW15-P*_fabA_* | This study |
| AK147 | MG1655 Δ*yhdP* Δ*fadR*::cm pJW15-P*_fabA_*(-19G>A) | This study |
| AK148 | MG1655 Δ*fabR* pJW15-P*_fabA_* | This study |
| AK149 | MG1655 Δ*fabR* pJW15-P*_fabA_*(-19G>A) | This study |

**Table S4. Primers used in this study**

| **Primer** | **Sequence (5ʹ to 3ʹ)** |
| --- | --- |
| K-*fadR*-FP (*fadR*::cm recombineering) | TCTGGTATGATGAGTCCAACTTTGTTTTGCTGTGTTATGGAAATCTCACTATGTGTAGGCTGGAGCTGCTTCG |
| K-*fadR*-RP (*fadR*::cm recombineering) | AACAACAAAAAACCCCTCGTTTGAGGGGTTTGCTCTTTAAACGGAAGGGACATATGAATATCCTCCTTAG |
| Overlap-pJW15-*fabA* FP | GGCCCTTTCGTCTTCACCTCGGAATCAGAGTATCGCTATCACAG |
| Overlap-pJW15-*fabA* RP | GATCCGGTACCCGGGCTGCAGTTCTCTGTAAGCCTTATTTTATTG |
| Overlap-pJW15-*fabA* FP1 | TGCAGCCCGGGTACCGGATCCTCTAGTTGCGGCCGCAAAATG |
| Overlap-pJW15-*fabA* RP1 | CGAGGTGAAGACGAAAGGGCCTCGTGATACGCCTATTTTTATAGG |

**Dataset 1 (separate file). Lipid extract LC-MS quantification.** Whole cell lipid extracts were subjected to LC/MS and absolute quantification for major phospholipids. Standard curves and measurements are shown for CL, PG, and PE. Peak areas for the analyte and internal standard are shown as well as the nmols of the indicated species per mg of protein.

**Dataset 2 (separate file). RNA-seq differentially regulated genes.** RNA-seq was performed with the wild-type, ∆*yhdP* ∆*fadR*, and Suppressor 1 strains. Differentially expressed genes were considered to have at least 2-fold change in expression and a p-value of <0.05 by quasi-linear F-test. Differentially expressed genes are shown for each strain comparison with log_2_ fold values, p-values, q-values (false discovery value), and expression values for each sample as normalized counts per million reads.

**Dataset 3 (separate file). All RNA-seq expression data.** RNA-seq was performed with the wild-type, ∆*yhdP* ∆*fadR*, and Suppressor 1 strains. Expression values as normalized counts per million reads are shown for all genes as well as log_2_ fold values, p-values, q-values for each strain pair.

**Dataset 4 (separate file). Raw data for all growth curves and reporter assays.**

Supplemental References

1.	Ruiz N, Davis RM, Kumar S. 2021. YhdP, TamB, and YdbH are redundant but essential for growth and lipid homeostasis of the gram-negative outer membrane. mBio 12:e02714-21.

2.	Mitchell AM, Srikumar T, Silhavy TJ. 2018. Cyclic enterobacterial common antigen maintains the outer membrane permeability barrier of *Escherichia coli* in a manner controlled by Yhdp. mBio 9:e01321-18.

3.	Rai AK, Carr JF, Bautista DE, Wang W, Mitchell AM. 2021. ElyC and cyclic enterobacterial common antigen regulate synthesis of phosphoglyceride-linked enterobacterial common antigen. mBio 12:e02846-21.

4.	Guyer MS, Reed RR, Steitz JA, Low KB. 1981. Identification of a sex-factor-affinity site in E. coli as gamma delta. Cold Spring Harb Symp Quant Biol 45 Pt 1:135-40.

5.	Mitchell AM, Wang W, Silhavy TJ. 2017. Novel Rpos-dependent mechanisms strengthen the envelope permeability barrier during stationary phase. J Bacteriol 199:e00708-16.

6.	Singer M, Baker TA, Schnitzler G, Deischel SM, Goel M, Dove W, Jaacks KJ, Grossman AD, Erickson JW, Gross CA. 1989. A collection of strains containing genetically linked alternating antibiotic resistance elements for genetic mapping of Escherichia coli. Microbiol Rev 53:1-24.

7.	Morris KN, Mitchell AM. 2023. Phosphatidylglycerol Is the Lipid Donor for Synthesis of Phospholipid-Linked Enterobacterial Common Antigen. Journal of Bacteriology 205:e00403-22.

8.	MacRitchie DM, Ward JD, Nevesinjac AZ, Raivio TL. 2008. Activation of the Cpx envelope stress response down-regulates expression of several locus of enterocyte effacement-encoded genes in enteropathogenic *Escherichia coli*. Infection and Immunity 76:1465-1475.
